# Co-surveillance of ribosomal RNA by the exosome complex and nucleolar RNAi in *C. elegans*

**DOI:** 10.1101/2020.11.23.395020

**Authors:** Shimiao Liao, Xiangyang Chen, Ting Xu, Qile Jin, Zongxiu Xu, Demin Xu, Xufei Zhou, Chengming Zhu, Shouhong Guang, Xuezhu Feng

**Affiliations:** Ministry of Education Key Laboratory for Membraneless Organelles & Cellular Dynamics, Hefei National Laboratory for Physical Sciences at the Microscale, School of Life Sciences, Division of Life Sciences and Medicine, University of Science and Technology of China, Hefei, Anhui 230027, P.R. China; CAS Center for Excellence in Molecular Cell Science, Chinese Academy of Sciences, Hefei, Anhui 230027, P.R. China

**Keywords:** nucleolus, rRNA, risiRNA, exosome, nucleolar RNAi, transcription

## Abstract

Eukaryotic cells express a wide variety of endogenous small regulatory RNAs that function in the nucleus. We previously found that erroneous rRNAs induce the generation of antisense ribosomal siRNAs (risiRNAs) which silence the expression of rRNAs via the nuclear RNAi defective (Nrde) pathway. To further understand the biological roles and mechanisms of this class of small regulatory RNAs, we conducted forward genetic screening to identify factors involved in risiRNA generation in *Caenorhabditis elegans*. We found that risiRNAs accumulated in the RNA exosome mutants. risiRNAs directed a NRDE-dependent silencing of pre-rRNAs in the nucleolus. In the presence of risiRNA, NRDE-2 accumulated in the nucleolus and colocalized with RNA polymerase I. risiRNA inhibited the transcription elongation of RNA polymerase I by decreasing RNAP I occupancy downstream of the site of RNAi. Meanwhile, exosome mislocalized from the nucleolus to nucleoplasm in suppressor of siRNA *(susi)* mutants, in which erroneous rRNAs accumulated. These results establish a novel model of rRNA surveillance by combining ribonuclease-mediated RNA degradation with small RNA-directed nucleolar RNAi system.

## Introduction

In eukaryotic cells, ribosomal RNAs (rRNAs) are transcribed by RNA polymerase I into a single 47S polycistronic precursor in the nucleolus, which are then processed and matured into 18S, 5.8S, and 28S rRNAs; 5S rRNA is independently transcribed by RNA polymerase III in the nucleus. The processing of ribosomal RNAs is extraordinarily complicated, in which defects of any steps could induce the accumulation of erroneous rRNAs [1, 2]. Immature rRNA intermediates or erroneous rRNAs are degraded by multiple surveillance machineries. In the nucleus, the RNA exosome has a central role in monitoring nearly every type of transcripts produced by RNA polymerase I, II, and III (RNAP I, II, and III) [3]. The eukaryotic nuclear RNA exosome is a 3’ to 5’ exoribonuclease complex, consisting of a 9-protein catalytically inactive core complex (EXO-9) and two catalytic subunits, Rrp6 (also known as EXOSC10), and Dis3 (also known as Rrp44 or EXOSC11)[4]. Erroneous rRNAs are degraded from 3’ to 5’ by the RNA exosome complex. In the cytoplasm, erroneous rRNAs can be polyuridylated and degraded from 3’ to 5’ by the cytoplasmic exoribonuclease DIS3L2 (also known as SUSI-1 in *C. elegans*) [5, 6].

rRNA-derived small RNAs have been identified in a number of organisms. In *Schizosaccharomyces pombe*, defects in TRAMP-mediated RNA surveillance system elicit the biogenesis of rRNA-siRNAs (rr-siRNAs) and reduce the levels of centromeric siRNAs [7]. In *Arabidopsis*, 24-or 21-nt rDNA-derived siRNAs have been identified and the latter siRNAs are accumulated upon viral infection or the depletion of the 5’ to 3’ RNA degradation machineries [8–12]. In *Neurospora crassa*, 20-to 21-nt qiRNAs are produced from aberrant rRNAs in an RNA-dependent RNA polymerase (RdRP)-dependent manner, and function in DNA damage repair [13]. In *C. elegans*, 22G antisense ribosomal siRNAs (risiRNAs) are generated upon environmental stresses or improper pre-rRNA processing [5, 14, 15]. risiRNAs down-regulate pre-rRNAs through the nuclear RNAi pathway in the nucleolus [16].

Small regulatory RNAs direct sequence-specific regulation of gene expression via the mechanism termed RNA interference (RNAi). Small RNAs guide the Argonaute-containing protein complex to complementary nucleic acids and modulate gene expression by a number of mechanisms, including but not limiting to RNA degradation, translation inhibition, inducing heterochromatin formation, and inhibiting transcription elongation [17, 18]. In *C. elegans*, siRNAs silence nuclear-localized RNAs co-transcriptionally via the Nrde pathway. The NRDE complex transports 22G siRNAs from the cytoplasm to the nucleus, inhibits RNA polymerase II during the elongation phase of transcription and induces histone H3 lysine 9 (H3K9) and histone H3 lysine 27 (H3K27) trimethylation [19–21]. Similarly, the nuclear Argonaute protein NRDE-3 bind risiRNAs and translocate from the cytoplasm to the nucleolus, in which the risiRNA/NRDE complex associates with pre-rRNAs and reduces the level of pre-rRNAs [5, 14]. However, the detailed mechanism of risiRNA-mediated pre-rRNA silencing is poorly understood.

To further understand the biological roles and mechanisms of risiRNA, in this study, we isolated a series of exosome mutants in which risiRNAs were accumulated by forward and reverse genetic screens and CRISPR-Cas9–mediated gene knockout technology. We found that the nucleolar localization of RNA exosome was important for risiRNA suppression. Meanwhile, we developed a RNAP I transcription activity assay and demonstrated that risiRNAs guide the NRDE complex to nucleoli to inhibit RNAP I transcription, a process independent of H3K9 and H3K27 trimethylation. Therefore, we concluded that cells combine ribonuclease-mediated RNA degradation with small RNA-directed nucleolar RNAi system to maintain rRNA homeostasis in *C. elegans*.

## Results

### Genetic screening identified risiRNAs that accumulated in the *susi-5(ceDis3)* mutant

We previously described a forward genetic screen used to search for suppressors of siRNA production in *C. elegans* (Fig. 1A) [5, 14]. This screen identified a cytoplasmic localized exoribonuclease SUSI-1(ceDIS3L2) and a number of rRNA modifying and processing enzymes (Fig. 1B). Here, we report that this screen identified a mutant allele, *ust56,* in *susi-5* gene that suppresses risiRNA production. In *eri-1(mg366);susi-5(ust56)* mutants, the argonaute protein NRDE-3 accumulates in the nucleus of seam cells in *C. elegans* (Fig. 1C). NRDE-3 transports siRNAs from the cytoplasm to the nucleus. NRDE-3 localizes to the nucleus when it binds to siRNAs but accumulates in the cytoplasm in the absence of siRNA ligands, as observed in the *eri-1* mutant [21]. The subcellular localization of NRDE-3 makes it a useful tool to monitor the abundance of cellular siRNAs [21]. The production of risiRNAs in *susi* mutants triggers the accumulation of NRDE-3 in the nucleus and nucleoli in an *eri-1*-independent manner [5, 14]. We deep sequenced small RNAs in control animals and the *susi-5(ust56)* mutant and observed an increase in risiRNAs (Fig. S1A).

**Fig. 1.**
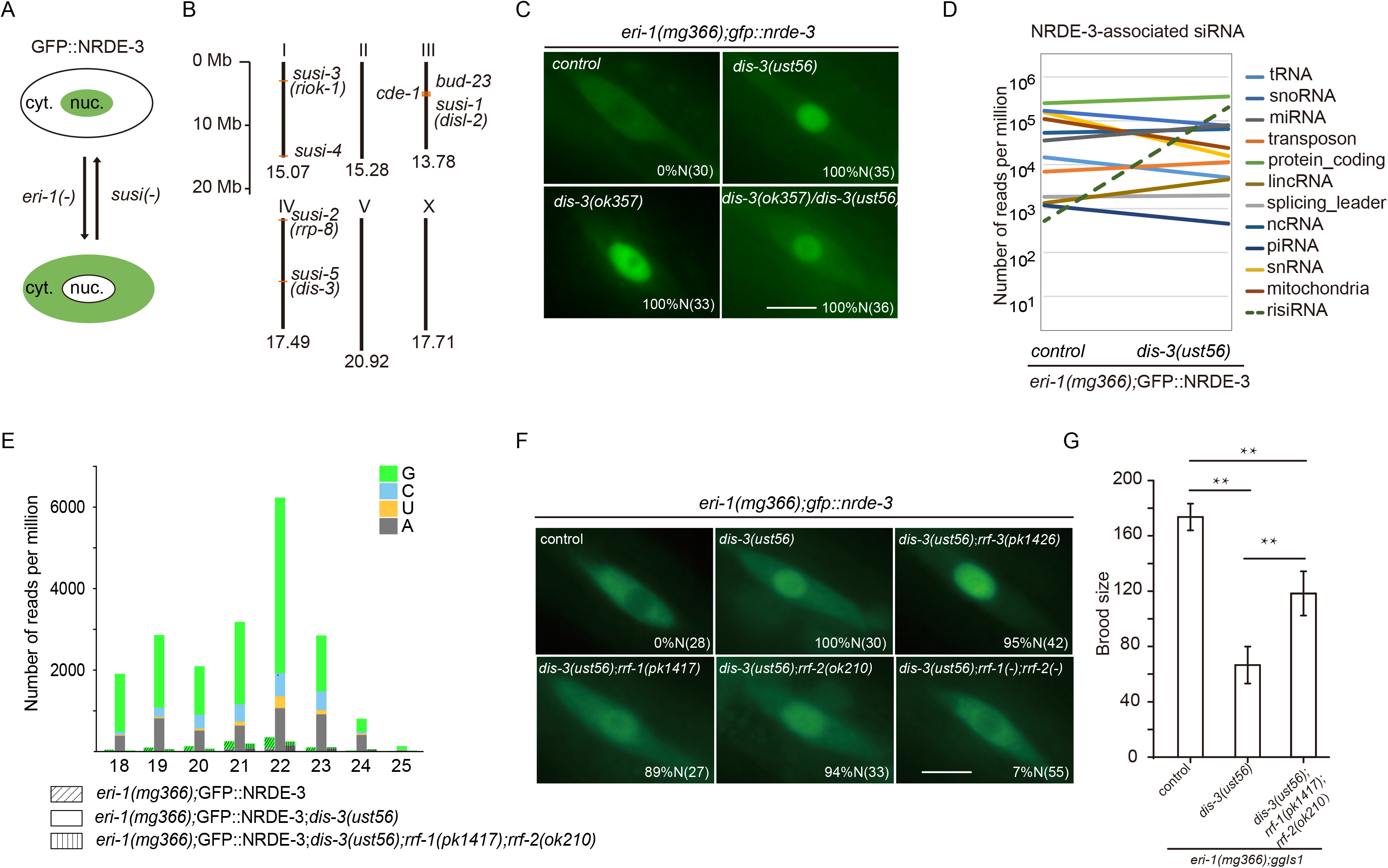
A genetic screen identified the accumulation of antisense ribosomal siRNA (risiRNA) in the *dis-3* mutant. (A) The subcellular localization of NRDE-3 was used as an indicator to screen for suppressors of mutants in endo-siRNA generation. cyt., cytoplasm; nuc., nucleus. (B) Summary of *susi* genes identified by forward genetic screening in *C. elegans* [5, 14]. Numbers indicate the size of each chromosome. (C) Images show seam cells of the indicated genotype expressing GFP::NRDE-3. Numbers indicate the percentage of animals with nucleus-enriched NRDE-3 in seam cells (% N). The number of scored animals is indicated in parentheses. Scale bars, 5 μm. (D) Results of the deep sequencing of NRDE-3-associated siRNAs from the indicated animals. The green dashed lines indicate risiRNAs. (E) Size distribution and 5’-end nucleotide preference of NRDE-3-associated risiRNAs in indicated animals. (F) Images show seam cells of the respective genotype expressing GFP::NRDE-3, labeled as in (C). Scale bars, 5 μm. (G) Brood size of indicated animals grown at 20°C. Data are presented as the mean ± s.d.; *n* ≥ 15 animals; **P < 0.01.

To determine the molecular identity of *susi-5, w*e mapped *susi-5(ust56)* to the open reading frame C04G2.6 by SNP mapping followed by genome resequencing. C04G2.6 is predicted to encode a protein that is homologous to yeast DIS3 and human RRP44 and engages in pre-rRNA surveillance [22]. C04G2.6 has a PIN domain, two cold shock domains (CSD), an RNB domain and S1 domain (Fig. S1B). While the CSDs and the S1 domains contribute to RNA binding, both the RNB and PIN domains are responsible for RNA degradation [23]. In the *ust56* allele, a conserved amino acid in the cold shock domain, Arg363, was mutated to cysteine. We acquired one additional allele, *c04g2.6(ok357)*, from the Caenorhabditis Genetics Center (CGC). NRDE-3 also accumulates in the nucleus of seam cells in the *eri-1(mg366);dis-3(ok357)* strain (Fig. 1C). An ectopically expressed mCherry::C04G2.6 transgene rescued the cytoplasmic localization of NRDE-3 in *eri-1(mg366);susi-5(ust56)* animals (Fig. S1C). Thus, we concluded that *susi-5* is *dis-3,* and the name *dis-3* is used hereafter. The *ok357* mutation deletes the PIN and two CSD domains and is a null allele (Fig. S1B). The *dis-3(ok357)* mutant is arrested at the L1 stage and has no progeny (Fig. S1D). However, the *dis-3(ust56)* strain is fertile, and approximately 180 progeny are produced per hermaphrodite at 20°C, suggesting that the R363C mutation partially disrupts DIS-3 function. In addition, *dis-3(ust56)* is a temperature sensitive allele. At 25°C, the *dis-3(ust56)* mutant is sterile. In this study, we used *dis-3(ust56)* as the reference allele.

To confirm that NRDE-3 associates with risiRNAs in *dis-3* mutants, we immunoprecipitated NRDE-3 from *dis-3(ust56)* mutant animals and deep sequenced NRDE-3-associated small RNAs. risiRNAs are enriched in *dis-3(ust56)* mutants compared to the level found in control strains (Figs. 1D and S2A). risiRNAs belong to the 22G-RNA category in *C. elegans*. The majority of risiRNAs start with a guanosine at the 5’-end and are 22 nt in length (Fig. 1E). The generation of risiRNAs requires two RNA-dependent RNA polymerases, RRF-1 and RRF-2, which are essential for the production of 22G-RNAs (Fig. S2B). In *rrf-1;rrf-2* double mutants, NRDE-3 binds substantially fewer risiRNAs (Figs. 1E and S2A) and accumulates in the cytoplasm (Fig. 1F). The presence of risiRNAs decreased the fertility of *C. elegans*. While the *dis-3(ust56)* mutation reduced the *eri-1(mg366);dis-3(ust56)* brood size, the *rrf-1* and *rrf-2* mutations partially restored the strain fecundity (Fig. 1G).

### risiRNA accumulated in the *exosome* mutants

DIS-3 is a core factor of the RNA exosome, which is a 3’ to 5’ exoribonuclease complex containing a 9-protein catalytically inactive core complex (EXO-9) and two catalytic active subunits, EXOS-10 and DIS-3 (Fig. 2A) [22]. EXO-9 forms a double-layered barrel-like structure that comprises six ribonuclease (RNase) pleckstrin homology (PH)-like proteins (EXOS-4.1, EXOS-4.2, CRN-5, EXOS-7, EXOS-8, and EXOS-9) and three S1/K homology (KH) “cap” proteins (EXOS-1, EXOS-2, and EXOS-3). All these factors are conserved from yeast to humans. Most of the RNA exosome subunits are essential, and loss-of-function mutations in them lead to larval development arrest or animal sterility at 20°C (Fig. S3A) [14]. To determine whether other components of the exosome complex, in addition to DIS-3, is involved in suppressing risiRNA production, we acquired mutants of the other exosome subunits, *exos-2(tm6653*), *exos-3(tm6844), exos-4.1(tm5568), exos-9(ok1635)* and *exos-10(ok2269)* from the National Bioresource Project and the CGC, and generated *exos-1(ust57)*, *exos-5(ust61), exos-7(ust62)* and *exos-8(ust60)* by CRISPR/Cas9-mediated gene deletion (Fig. S3B-J). In all of the mutants, NRDE-3 accumulates in the nucleus in seam cells (Fig. 2B). We deep sequenced total small RNAs and NRDE-3-associated small RNAs in control animals and *eri-1(mg366);exos-1(ust57)* and *eri-1(mg366);exos-10(ok2269)* mutants and observed an increase in both total risiRNAs and NRDE-3-associated risiRNAs (Fig. 2C-D). Thus, we concluded that the exosome complex is involved in the suppression of risiRNA production.

**Fig. 2.**
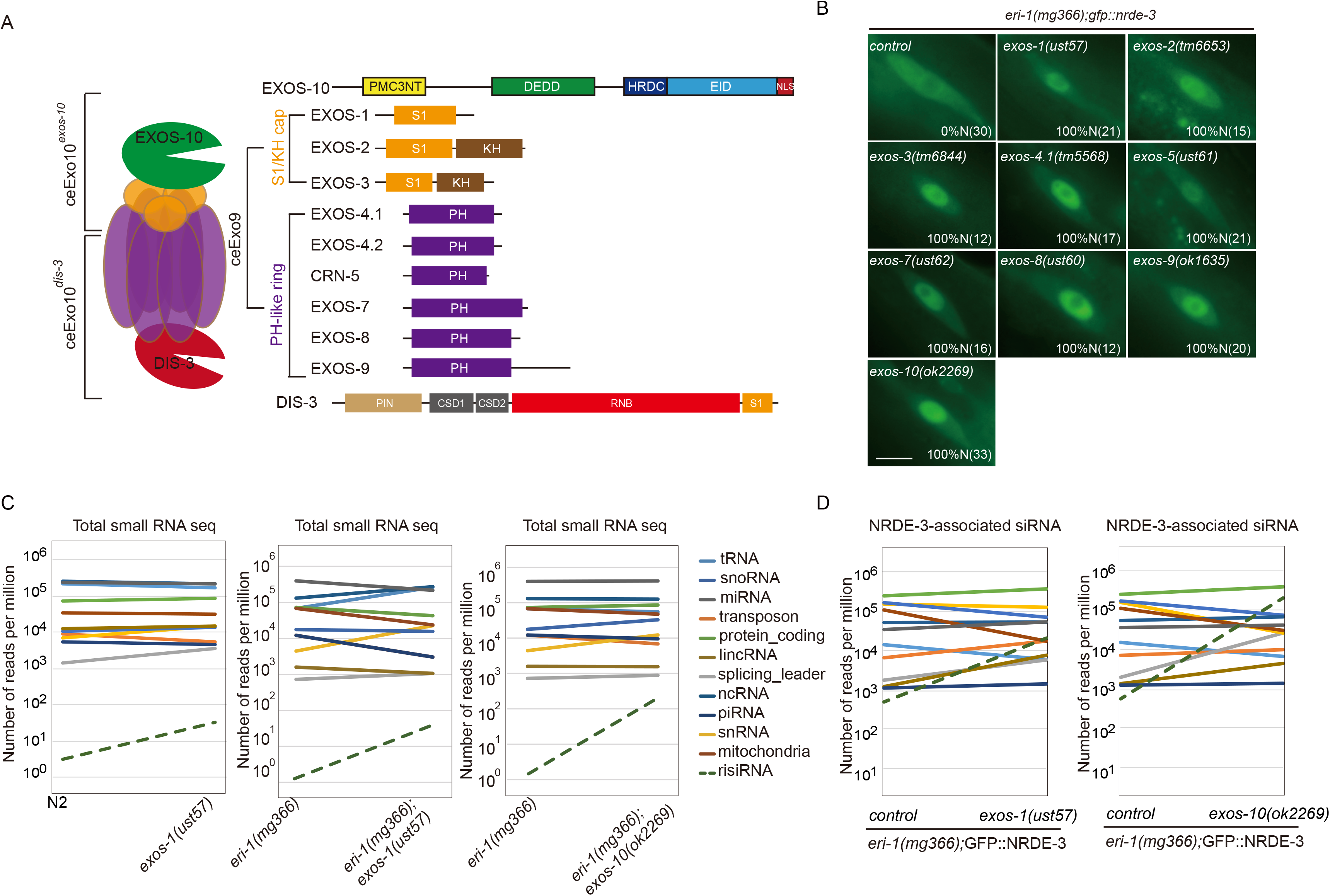
risiRNAs are enriched in exosome mutants. (A) Schematics of the RNA exosome complex and the subunits in *C. elegans.* (B) Images show seam cells of the respective genotype expressing GFP::NRDE-3. Numbers indicate the percentage of animals with nucleus-enriched NRDE-3 in seam cells (% N). The number of scored animals is indicated in parentheses. Scale bars, 5 μm. Schematics of the alleles are shown in Fig. S3. (C) Results from the deep sequencing of total small RNAs from the indicated animals. The green dashed lines indicate risiRNAs. (D) Results from the deep sequencing of NRDE-3-associated siRNAs from the indicated animals.

### risiRNA inhibites the RNA polymerase I-directed transcription

We further investigated the molecular mechanism of nucleolar RNAi. Small interference RNAs guide the NRDE complex to targeted pre-mRNAs, induce H3K9, H3K23 and H3K27 trimethylation at the corresponding genomic loci, inhibit RNAP II-mediated transcription elongation, and silence gene expression in the nucleus in *C. elegans* [19, 20, 24, 25]. To determine whether risiRNAs induce histone modifications, we conducted ChIP assays with anti-H3K9me3 and anti-H3K27me3 antibodies. However, we failed to identify a significant change in H3K9 and H3K27 trimethylation at the rDNA loci in the presence of risiRNAs (Fig. 3A). As a positive control, dsRNAs targeting the *lin-15b* gene, encoding an RNAP II transcript, induced both H3K9 and H3K27 trimethylation, as reported previously [20, 24].

**Fig. 3.**
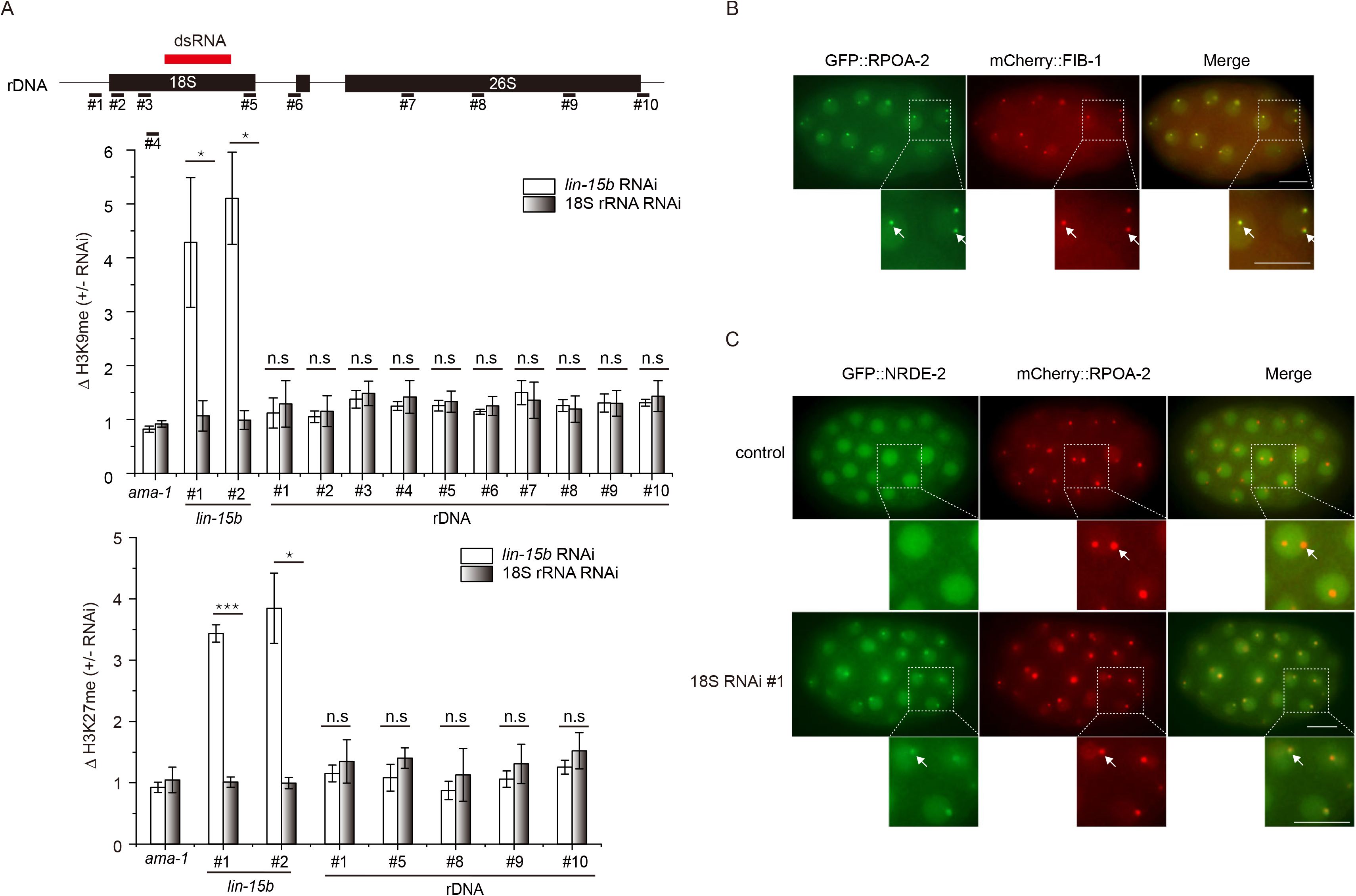
risiRNA directs NRDE-2 to the transcription site of rRNAs. (A) ChIP analysis of rDNA loci upon treatment of RNAi targeting the *lin-15b* gene or 18S rRNA. (top) Schematic of the rDNA transcription unit and real time PCR primers. The red bar indicates the dsRNA segment targeting 18S rRNA. Trimethylation of H3K9 and H3K27 (bottom) were measured by ChIP assay. Data are presented as mean ± s.d.; n = 3. *p<0.05, ***p<0.0001, n.s., not significant. (B) Images of *C. elegans* embryos expressing GFP:RPOA-2 (green) and mCherry::FIB-1 (red). (C) Images of *C. elegans* embryos expressing GFP:NRDE-2 (green) and mCherry::RPOA-2 (red) after the animals were treated with RNAi targeting 18S rRNA. Scale bars, 10 μm.

To determine whether risiRNA-guided nucleolar RNAi silence rRNAs by inhibiting RNAP I-directed transcription elongation, we first generated GFP- and mCherry-tagged RPOA-2 transgene *in situ* by CRISPR/Cas9 technology. RPOA-2 is the core subunit of RNAP I and contribute to polymerase activity [26]. Knocking down RPOA-2 by RNAi caused sterility in the animals, suggesting that RPOA-2 play essential roles (Fig. S4A). RPOA-2 was enriched in the nucleoli and colocalized with the nucleoli marker FIB-1 (Figs. 3B and S4B-C), a finding that is consist with their functions in rRNA transcription. In 1-to 8-cell embryos, in which FIB-1 foci is absent, RPOA-2 was evenly distributed in the nucleus without significant nucleolar enrichment (Figs. S4C), a finding that is consistent with the idea that rDNA is not actively transcribed in early embryos. NRDE-2 is ubiquitously expressed and evenly distributed in the nucleus in *C. elegans* [19]. In the presence of risiRNAs, NRDE-2 was enriched in the nucleoli and colocalized with RPOA-2 (Figs. 3C and S4D), suggesting that risiRNA guides NRDE-2 to pre-rRNA and modulates rRNA transcription.

To further determine how risiRNAs silence rRNA expression, we validated the activity of GFP::RPOA-2 by ChIP assay. Actinomycin D is able to block the transcriptional activity of both RNAP I and RNAP II. When animals were treated with actinomycin D, RPOA-2 paused at the 5′-end of the rRNA transcription unit and failed to elongate toward the 3′-end (Fig. 4A), suggesting that the GFP::RPOA-2 fusion protein recapitulates the function of endogenous proteins. Actinomycin D treatment did not substantially change the expression level and subcellular location of RPOA-2, FIB-1, EXOS-10 and NRDE-3 (Figs. S5A-C and S6A-B).

**Fig. 4.**
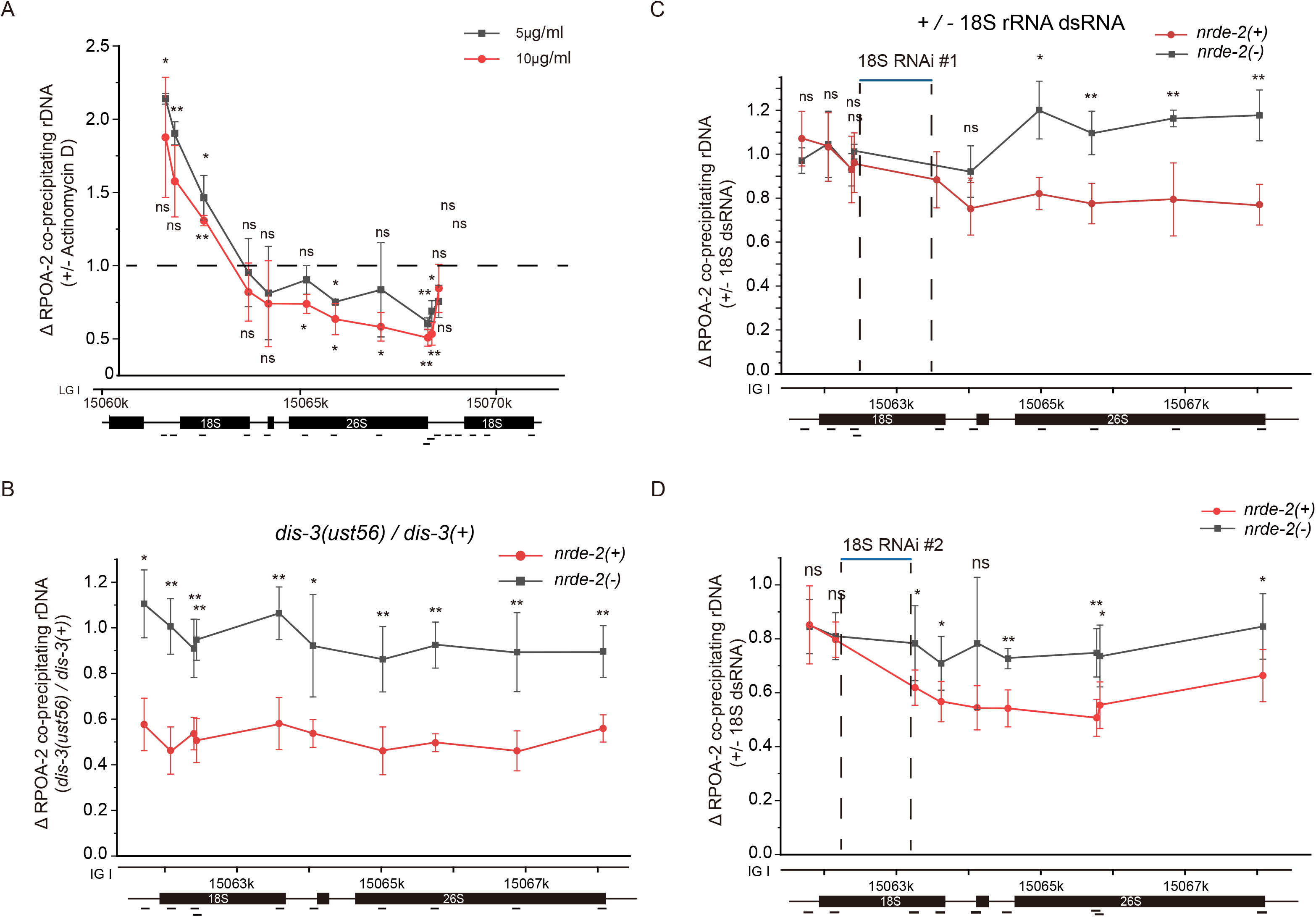
risiRNA directs a NRDE-2-dependent inhibition of RNAP I transcription. (A) ChIP assay of RPOA-2 occupancy upon actinomycin D treatment. Fold changes were normalized to 1% input first, and then compared to the no-drug treatment group. Mean ± s.d.; n = 4; *P<0.05 and **P<0.01. (B) Results of the ChIP assay of RPOA-2 occupancy in the indicated animals. The enrichment of each sample was first normalized to 1% input. And then fold changes were calculated by dividing the enrichment of *dis-3(ust56)* mutants by the number of control animals. Statistics were analyzed by comparing the data from the *nrde-2(+)* and *nrde-2(-)* animals. mean ± s.d.; n = 4; *P<0.05, **P<0.01. (C, D) RPOA-2 occupancy along the rDNA unit was quantified by ChIP-qPCR upon RNAi targeting of 18S rRNA regions to indicated animals. Statistics were analyzed by comparing the data obtained for the *nrde-2(+)* and *nrde-2(-)* animals; mean ± s.d.; n = 5; *P < 0.05 and ** P< 0.01.

We then investigated whether risiRNA silences rRNA expression by inhibiting RNAP I transcription elongation. We examined GFP::RPOA-2 occupancy by ChIP assay of the control animals, *dis-3* mutants, and animals being treated with RNAi targeting a fragment of 18S rRNA. In *dis-3(ust56)* mutants, GFP::RPOA-2-associated rDNA was pronouncedly decreased, a phenomenon dependent on the nuclear RNAi factor NRDE-2 (Fig. 4B). In the absence of *nrde-2*, no change in RPOA-2 occupancy was observed. Treating animals with exogenous RNAi targeting 18S rRNA had no significant effect on RPOA-2 occupancy near the site of transcription initiation and upstream of the RNAi-targeted site (Figs. 4C-D). However, we detected a decrease in RPOA-2 occupancy downstream of the RNAi-targeted region. In addition, in the absence of *nrde-2*, risiRNAs failed to reduce RPOA-2 occupancy downstream of the RNAi-targeted region. Similar inhibition of transcription elongation had been observed for RNA polymerase II transcripts during nuclear RNAi [19]. Taken together, these data suggest that risiRNAs, acting together with the nuclear RNAi machinery, may silence nascent RNAP I transcripts during the elongation phase of transcription in *C. elegans*.

### The nucleolar localization of exosomes was important for risiRNA suppression

To further investigate the biological roles of the exosome complex in risiRNA generation in *C. elegans*, we constructed fluorescent protein-tagged exosome subunits, including mCherry::DIS-3, GFP::EXOS-1, GFP::EXOS-2, and GFP::EXOS-10. These subunits are ubiquitously expressed in all cells and enriched in the nucleus (Figs. 5A and S7A-B). We also constructed mCherry- and GFP-tagged SUSI-2(ceRRP8), a protein previously identified as a suppressor of risiRNA production [14]. *C. elegans* SUSI-2 is the homolog of yeast protein RRP8, which exclusively localizes in the nucleolus and engages in m1A methylation of the 26S rRNA. We crossed GFP::EXOS-1 and GFP::EXOS-10 onto a mCherry::SUSI-2 background, respectively, and found that EXOS-1 and EXOS-10 were enriched in nucleoli and colocalized with SUSI-2 in somatic cells (Figs. 5B and S8A). After crossing mCherry::DIS-3 with GFP::SUSI-2 animals, we found that DIS-3 was depleted from nucleoli but enriched in the nucleoplasm (Figs. 5C and S8B). We crossed GFP::EXOS-1 and GFP::EXOS-10 with *dis-3(ust56)* mutants. Surprisingly, EXOS-1 and EXOS-10 were depleted from the nucleoli but enriched in the nucleoplasm (Fig. 5D).

**Fig. 5.**
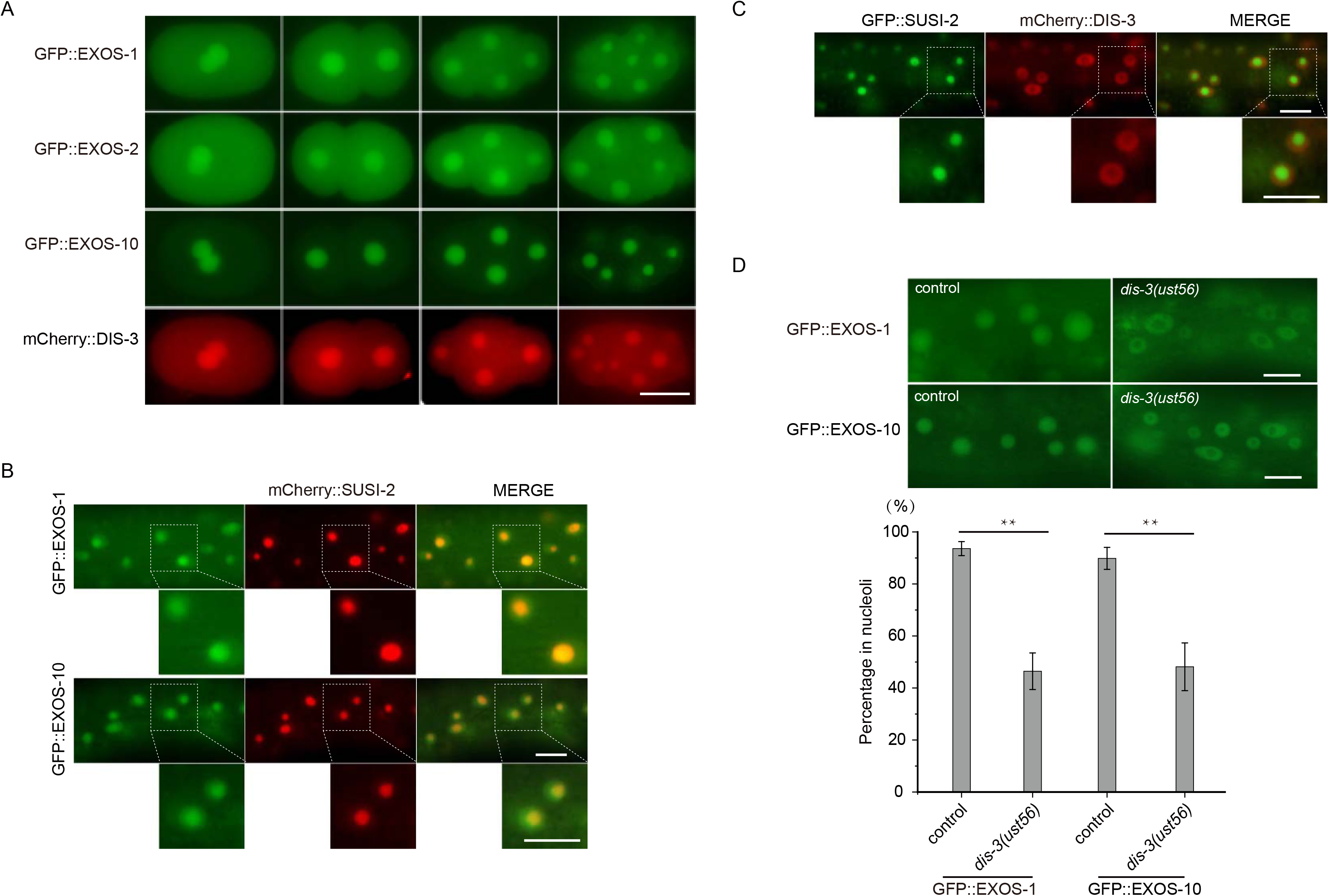
*dis-3(ust56)* mutation mislocalized exosome components from the nucleoli to nucleoplasm. (A) Images of 1-to-8 cell embryos from the animals expressing indicated transgenes. Scale bars, 10 μm. (B) Images show somatic cells of animals expressing GFP::EXOS-1 (green), GFP::EXOS-10 (green) and mCherry::SUSI-2 (red). Animals were grown at 20°C. Scale bars, 10 μm. (C) Images show somatic cells of the animals expressing GFP::SUSI-2 (green) and mCherry::DIS-3 (red). Scale bars, 10 μm. (D) (top) Images of somatic cells of the indicated animals. Scale bars, 10 μm. (bottom) Quantification of the nucleolar localization of GFP::EXOS-1 and GFP::EXOS-10. mean ± s.d.; n > 70 animals; **P < 0.01.

Two lines of evidence suggested that the nucleolar localization of exosomes is important for risiRNA suppression. First, *susi-2(ceRRP8)* and T22H9.1 are known SUSI proteins and are involved in the modification and processing of rRNAs [14]. Knocking down *susi-2(ceRRP8)* and T22H9.1 by RNAi induced increase of risiRNAs [14] and depletion of GFP::EXOS-10 from the nucleoli (Fig. 6A). Second, we performed a candidate-based RNAi screen to search for rRNA processing factors that are required for the nucleolar localization of GFP::EXOS-10. We selected fifteen predicted rRNA processing factors and investigated whether knocking down these genes by RNAi could block the nucleolar accumulation of EXOS-10 (Table S1). We found that knocking down *M28.5*, *nol-56*, *fib-1* and *mtr-4* by RNAi induced the depletion of EXOS-10 from the nucleoli (Fig. 6A). Among the proteins encoded by these genes, NOL-56 is an ortholog of human NOP56, which binds snoRNAs and facilitates box C/D ribonucleoprotein-guided methyltransferase activity [27]. FIB-1 encodes the *C. elegans* ortholog of human fibrillarin and *Saccharomyces cerevisiae* Nop1p. FIB-1 has RNA binding and rRNA methyltransferase activities, which are essential for nucleologenesis [28]. MTR-4 is an ortholog of human MTREX and has ATP-dependent RNA helicase activity [22]. We further deleted *fib-1*, *nol-56* and *mtr-4* by CRISPR/Cas9 technology (Fig. S9A). In these mutants, NRDE-3 redistributed from the cytoplasm to the nucleus in seam cells (Fig. 6B). In addition, after knocking down these genes by RNAi, risiRNAs were enriched, as shown by small RNA deep sequencing (Fig. 6C).

**Fig. 6.**
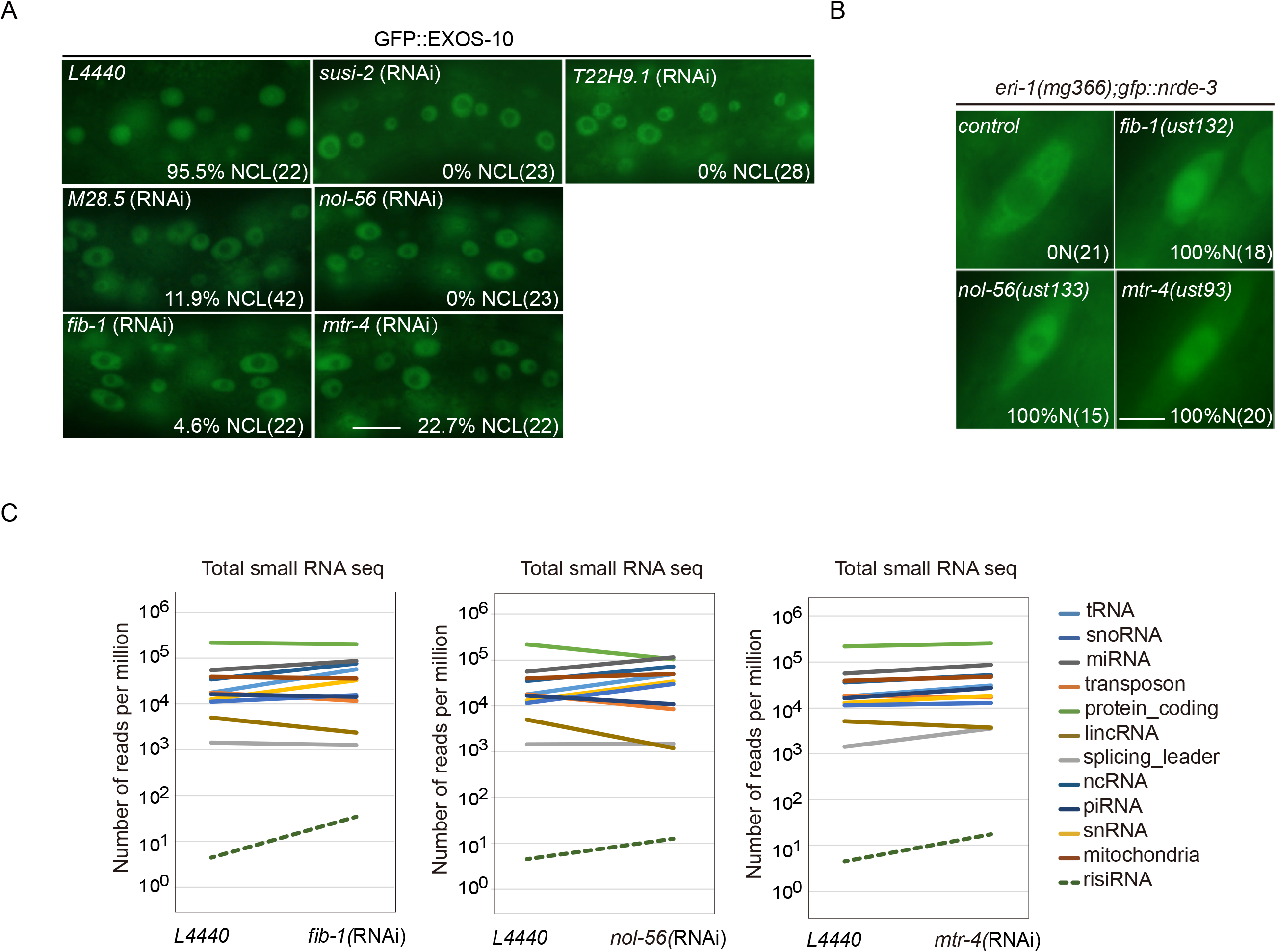
A proper nucleolar localization of the exosome complex is important for risiRNA suppression. (A) Images of somatic cells of the animals expressing GFP::EXOS-10 after being treated with RNAi targeting the indicated genes. The percentage of animals with nucleolar localized GFP::EXOS-10 is indicated (% NCL). The number of scored animals is indicated in parentheses. Scale bars, 10 μm. Animals were grown at 20°C. (B) Images of seam cells from the indicated animals. Numbers indicate the percentages of the animals with nuclear-enriched NRDE-3 in seam cells (%N). The number of scored animals is indicated in parentheses. Scale bars, 5 μm. (C) Results from the deep sequencing of total small RNAs from the indicated animals.

These data suggest that proper nucleolar localization of the exosome complex may be important for the suppression of risiRNA production, and can be used as a tool to screen for new *susi* genes. Yet a direct causative relationship between exosome mislocalization and risiRNA production remains to be determined.

## Discussion

Eukaryotic cells express a multitude of small regulatory RNAs and antisense transcripts that are of unknown function [29]. Small RNAs have been shown to induce endonucleolytic cleavage of target RNAs (slicer activity) or induce epigenetic modifications, including DNA and histone modifications [30]. In *C. elegans*, the nuclear argonaute protein NRDE-3 in soma (or HRDE-1 in the germline) lacks the residues required for slicer activity but inhibits RNAP II-mediated transcription elongation in the presence of siRNAs [19]. Here, we showed that risiRNAs guide the NRDE complex to pre-rRNAs to inhibit RNAP I transcription, a process independent of H3K9 and H3K27 trimethylation (Fig. 7). Thus, our data suggest a mechanism for nucleolar RNAi: risiRNA-directed cotranscriptional silencing of RNAP I. NRDE-2 is a conserved protein, which is involved in processing of pre-mRNAs in mammalian cells [31–33]. It will be of interest to investigate whether risiRNAs and RNAP I are similarly linked in other metazoans [12].

**Fig. 7.**
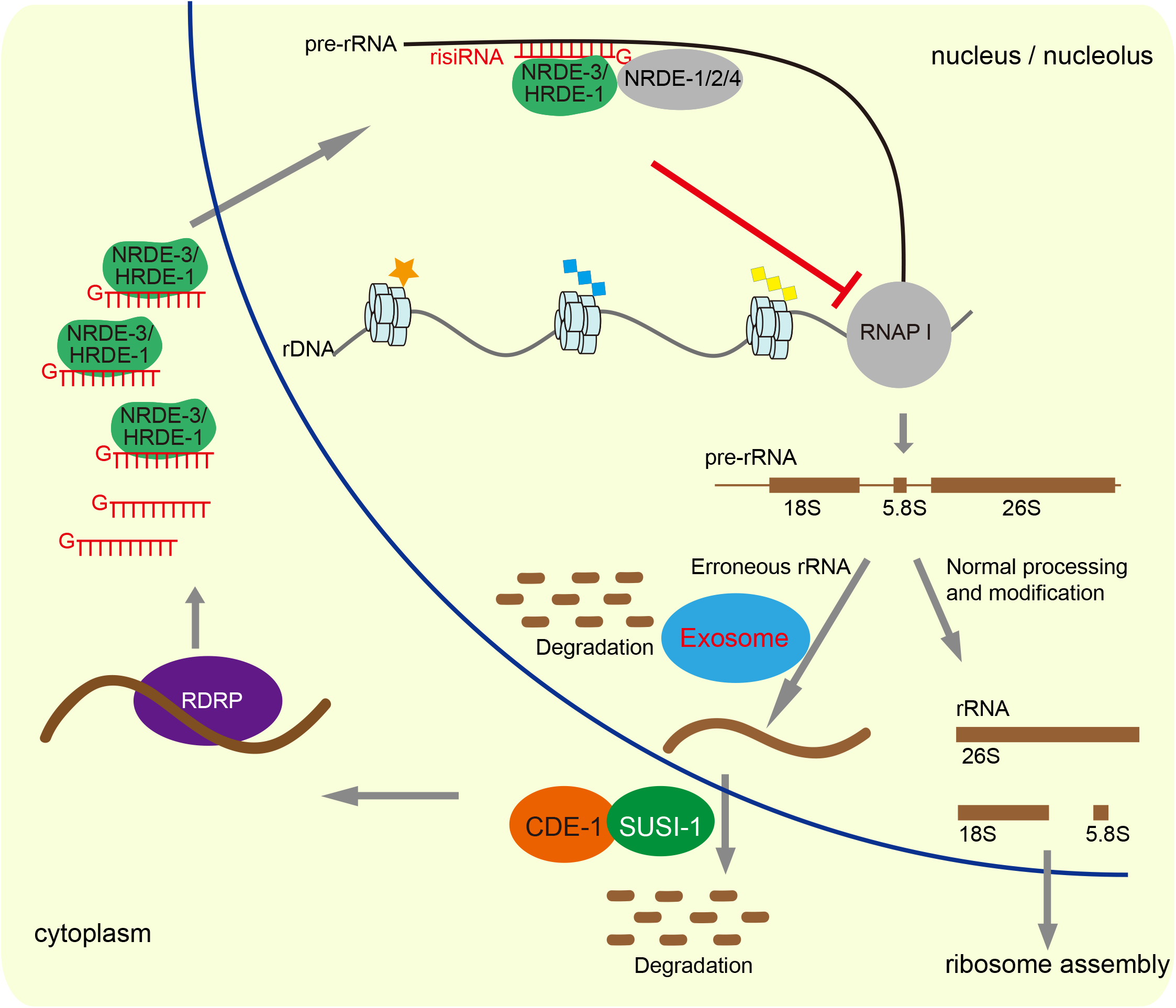
A working model of risiRNA biogenesis and function. The processes of ribosome biogenesis are very sophisticated in eukaryotic cells from the splicing events of pre-rRNAs to the final assemblage of ribosomes, during which errors could occur at any step. In the nucleoli and nucleus, an exoribonucleolytic multisubunit protein complex, the exosome, participates in rRNA processing and degradation. In the cytoplasm, erroneous rRNAs are uridylated at the 3’-ends by polyuridylating polymerase-I (named CDE-1 or PUP-1) and then degraded by the 3’ to 5’ exoribonuclease SUSI-1(ceDIS3L2). Deficiency of these two degradation systems results in the accumulation of erroneous uridylated rRNAs, which further recruit additional RNA-dependent RNA polymerases (RdRPs) to synthesize risiRNAs and initiate the nucleolar gene silencing cascade. risiRNAs associate with the nuclear argonaute protein NRDE-3 in soma or HRDE-1 in the germline, bind to pre-rRNAs, and inhibit RNAP I transcription elongation. Therefore, by combining the RNA degradation system with nucleolar gene silencing machinery, cells surveil the quality of rRNAs and maintain the homeostasis.

Small-RNA-guided chromatin modifications have been widely studied in many organisms. In *C. elegan*s, NRDE complex transports 22G RNAs from the cytoplasm to the nucleus, induces H3K9, H3K23 and H3K27 trimethylation and mediates transgenerational inheritance of RNAi [25, 34]. In this study, we found that the risiRNA/NRDE complex inhibits RNAP I transcription without altering the status of H3K9 and H3K27 trimethylation of rDNA genes. rDNA is a multicopy gene while only a proportion of the copies are actively transcribed in many organisms [35–38]. In *S. cerevisiae*, actively transcribed rDNA genes are largely devoid of histone molecules and are organized in a specialized chromatin structure that binds the high-mobility group protein Hmo1 [39]. Reducing the rDNA transcription efficiency upon the depletion of *dao-5* does not induce significant change of H3K9me3 modification at rDNA region in *C. elegans* [40]. Further study to identify which types of histone modifications are engaged in rDNA silencing will facilitate the understanding of the mechanism and regulation of risiRNA-directed RNAP I inhibition.

The processing of ribosomal RNAs is extraordinarily complicated and defects may occur at every step from production to assembly and cause ribosomopathies [1, 41]. Multiple surveillance machineries, including the nuclear-localized RNA exosome complex and the cytoplasmic exoribonuclease SUSI-1(ceDIS3L2), degrade defective rRNAs [3, 22, 42, 43]. Surveillance machinery deficiencies result in the accumulation of erroneous rRNAs. However, *C. elegans* utilizes a backup system, nucleolar RNAi, in which risiRNAs are produced to induce a nucleolar gene silencing by inhibiting RNAP I transcription. Therefore, these two systems act together to maintain rRNA homeostasis and prohibit the accumulation of erroneous rRNAs.

## Materials and methods

### Strains

Bristol strain N2 was used as the standard wild-type strain. All strains were grown at 20°C unless specified. The strains used in this study are listed in Supplemental Table S2.

### Genetic screening

Genetic screen experiment was conducted as previously described [5]. Briefly, to identify the factors which negatively regulate endo-siRNA generation, we searched for mutants that redistributed NRDE-3 from the cytoplasm to the nucleus in *eri-1(mg366);gfp::nrde-3* animals. NRDE-3 transports siRNAs from the cytoplasm to the nucleus. NRDE-3 localizes to the nucleus when it binds to siRNAs but accumulates in the cytoplasm in the absence of siRNA ligands, for example, in the *eri-1* mutant [21]. The production of risiRNAs in *susi* mutants triggers the accumulation of NRDE-3 in the nucleus and nucleoli. *eri-1(mg366);gfp::nrde-3* animals were mutagenized by ethyl methanesulfonate (EMS), followed by a clonal screen. The F2 progeny worms were visualized under fluorescence microscope at the L3/L4 stage. Mutants that redistributed NRDE-3 to the nuclei of seam cells were selected. *susi-5* was identified by snp-SNP mapping followed by the re-sequencing of the mutant genome.

### Construction of plasmids and transgenic strains

For in situ transgene *3xflag::gfp::rpoa-2*, the 3xFLAG::GFP coding region was PCR amplified from YY174 genomic DNA with the primers 5′-ATGGACTACAAAGACCATGACGG-3′ and 5′-AGCTCCACCTCCACCTCCTTTGTATAGTTCATCCATGCCATGT-3′. A 1.5kb homologous left arm was PCR amplified with the primers 5′-GGGTAACGCCAGCACGTGTGGTCAATGTCTAACAGCCAGCGAC-3′ and 5′-TCATGGTCTTTGTAGTCCATTATGTCGCAGTCCATCGCCTGA-3′. A 1.5kb homologous right arm was PCR amplified with the primers 5′-AAGGAGGTGGAGGTGGAGCTATGGACTGCGACATAGCGTCG-3′ and 5′-GAGTGAGCTGATACCAGCGGATGTACTTTGGCAACTTTAACAAATTG-3′. And the backbone was PCR amplified from the plasmid pCFJ151 with the primers 5′-CACACGTGCTGGCGTTACCC-3′ and 5′-CCGCTGGTATCAGCTCACTCAA-3′. All these fragments were joined together by Gibson assembly to form the *3xflag::gfp::rpoa-2* plasmid with the ClonExpress MultiS One Step Cloning Kit (Vazyme Biotech, Nanjing, China, Cat. No. C113-01/02). This plasmid was co-injected into N2 animals with three sgRNA expression vectors, rpoa-2_sgRNA#1, rpoa-2_sgRNA#2, rpoa-2_sgRNA#3, 5 ng/μl pCFJ90 and Cas9 II expressing plasmid. Primer pairs for constructing sgRNA expression vectors are shown in Table S3.

For in-situ transgene *mCherry::rpoa-2*, the mCherry fragment was amplified with the primers 5′-ATGGTCTCAAAGGGTGAAGAAG-3′ and 5′-ATAGCTCCACCTCCACCTCCCTTATACAATTCATCCATGCCACC-3′ and the vector plasmid was amplified with the primers 5′-GGAGGTGGAGGTGGAGCTATGGACTGCGACATAGCGTC-3′ from the *gfp::rpoa-2* plasmid. The two fragments were joined together by Gibson assembly to form the *mCherry::rpoa-2* repair plasmid. CRISPR plasmid mixture containing 30ng/μl rpoa-2_sgRNA#1, 30ng/μl rpoa-2_sgRNA#2, 30ng/μl rpoa-2_sgRNA#3, 50 ng/μl Cas9 II expressing plasmid and 5 ng/μl pCFJ90 was co-injected into N2 animals.

For in-situ transgene *3xflag::gfp::nrde-2*, the *3xflag* fragment was amplified with the primers 5′-ATGGACTACAAAGACCATGAC-3′ and 5′-ATAGCTCCACCTCCACCTCCTTTGTATAGTTCATCCATGCC-3′ from YY174 genomic DNA. A 1.5kb homologous left arm was PCR amplified with the primers 5′-GGGTAACGCCAGCACGTGTGGTCAATGTCTAACAGCCAGCGAC-3′ and 5′-TCATGGTCTTTGTAGTCCATATACGCTCGAAACATTGTTCATTA-3′. A 1.5kb homologous right arm was PCR amplified with the primers 5 ′-GGAGGTGGAGGTGGAGCTATGTTTCGAGCGTATGGAAATAATG-3′ and 5′-GCGGATAACAATTTCACCTAGATTATCCGAATCGTTTGCTAGAAC-3 ′. The backbone was PCR amplified with the primers 5 ′-TAGGTGAAATTGTTATCCGCTGG-3 ′ and 5 ′-TATTTCACACCGCATATGGTGC-3 ′ from pCFJ151. All these fragments were joined together by Gibson assembly to form the 3xflag::gfp::nrde-2 plasmid. This plasmid was co-injected into N2 animals with two sgRNA expression vectors, nrde-2_sgRNA#1, nrde-2_sgRNA#2 and Cas9 II expressing plasmid. Primer pairs for constructing sgRNA expression vectors are shown in Supplemental Table S3.

For the constructing of *mcherry::dis-3*, a 2kb promoter region was amplified with the primers 5′-CGACTCACTAGTGGGCAGATATCGTCGTGATTATCCATTTTTGAAAC-3′ and 5′-TCTTCACCCTTTGAGACCATGACGTTCAAATCCATACCTTC′. The *dis-3* CDs region and 3′ UTR region were amplified as a whole fragment with the primers 5′-GGAGGTGGAGGTGGAGCTATGGATTTGAACGTCAAACAAAG-3′ and 5′-GGCCTTGACTAGAGGGTACCAGCCGTCCCTATTGGATGATAAAT-3′. The *mCherry* coding sequence was amplified from PFCJ90 with 5′-AGCTCCACCTCCACCTCCCTTATACAATTCATCCATGCC-3′ and 5′-ATGGATTTGAACGTCATGGTCTCAAAGGGTGAAGAAGA-3′. The linearized backbone was amplified from PCFJ151 with primers 5′-ATCTGCCCACTAGTGAGTCG-3′ and 5′-GGTACCCTCTAGTCAAGGCC-3′. The transgene was integrated onto the *C. elegans*’ chromosome III of the strain EG8080 by MosSCI technology [44].

For *3xflag::gfp::exos-1*, a 2kb promoter region was amplified with the primers 5′-CGACTCACTAGTGGGCAGATTGCCTGACCTTAAGGCGG-3′ and 5′-TCATGGTCTTTGTAGTCCATCGTTTCGGCGAGCATTTTCT-3′. The *exos-1* CDs region and 3′ UTR region was amplified as a whole fragment with the primers 5′-AAGGAGGTGGAGGTGGAGCTATGCTCGCCGAAACGCTTGT-3′ and 5′-GGCCTTGACTAGAGGGTACCCAGTGAGCCCATCTCATCAT-3′. The *3xflag::gfp* coding sequence was amplified from YY174 genomic DNA with 5′-ATGCTCGCCGAAACGATGGACTACAAAGACCATGACGGTG-3′ and 5′-AGCTCCACCTCCACCTCCTTTGTATAGTTCATCCATGC-3′. The linearized backbone was amplified from pCFJ151 with primers 5′-ATCTGCCCACTAGTGAGTCG-3′ and 5′-GGTACCCTCTAGTCAAGGCC-3′. The transgene was integrated onto the *C. elegans*’ chromosome II of the strain EG4322 by MosSCI technology.

For *3xflag::gfp::exos-2*, a 2kb promoter region was amplified with the primers 5′-CGACTCACTAGTGGGCAGATACGAGAACAATCAAAGCAACG-3′ and 5′-TCATGGTCTTTGTAGTCCATGGTGACTTCGAAACTCATTT-3′. The *exos-2* CDS region and 3′ UTR region were amplified as a whole fragment with the primers 5′-AAGGAGGTGGAGGTGGAGCTATGAGTTTCGAAGTCACCGG-3′ and 5′-GGCCTTGACTAGAGGGTACCCGGTACCAACAACTCCAACG-3′. The *3xflag::gfp* coding sequence was amplified from YY174 genomic DNA with 5′-ATGGACTACAAAGACCATGACG-3′ and 5′-AGCTCCACCTCCACCTCCTTTGTATAGTTCATCCATGCCA-3′. The linearized backbone was amplified from pCFJ151 with primers 5′-ATCTGCCCACTAGTGAGTCG-3′ and 5′-GGTACCCTCTAGTCAAGGCC-3′. The transgene was integrated onto the *C. elegans*’ chromosome II of the strain EG4322 by MosSCI technology.

*exos-10* locates in the operon CEOP2496. For the constructing of *3xflag::gfp::exos-10,* a 2.1kb promoter region was PCR amplified with the primers 5′-CGACTCACTAGTGGGCAGATCAACGTCGGACTTCTCGAAT-3′ and 5′-CATATCTTGATAATCGTCCTCAT-3′ from N2 genomic DNA. A transpliced sequence was amplified with the primers 5′-AGGACGATTATCAAGATATGATGACGACATGCACTTTATA-3′ and 5′-TTCTTCTCCTGACATTCTGTAAAT-3′. The 3xFLAG::GFP coding region was PCR amplified from YY174 genomic DNA with the primers 5′-ACAGAATGTCAGGAGAAGAAGACTACAAAGACCATGACGGT-3′ and 5′-ATTGATTCTTCTCCTGACATAGCTCCACCTCCACCTCCT-3′. The EXOS-10 coding region and 3’ UTR region were PCR amplified with the primers 5′-ATGTCAGGAGAAGAATCAATGC-3′ and 5′-GGCCTTGACTAGAGGGTACCTGGATCTGAAGCTTAACCTATTC-3′. The pCFJ151 vector fragment was PCR amplified with the primers 5′-GGTACCCTCTAGTCAAGGCC-3′ and 5′-ATCTGCCCACTAGTGAGTCG-3′ from the pCFJ151 plasmid. These five fragments were joined together by Gibson assembly to form the *gfp::exos-10* repair plasmid. The transgene was integrated onto the *C. elegans*’ chromosome II by MosSCI technology.

For *susi-2::mCherry*, the promoter region was PCR amplified with the primers 5′-CCTGTCAATTCCCAAAATACTTGGAAAGCATTTTCAGGCG-3′ and 5′-GAAAAATTCAACGGAATGCTCTGAAATTGTTAACACAGATGATAAAAG-3′ and the coding region was PCR amplified with the primers 5′-AGCATTCCGTTGAATTTTTCGCTG-3′ and 5′-CAGCTCCACCTCCACCTCCGCGTTTCTTATACAAACAAGGC-3′ from N2 genomic DNA respectively. The mCherry fragment was PCR amplified with the primers 5′-CGGAGGTGGAGGTGGAGCTGTCTCAAAGGGTGAAGAAGATAAC-3′ and 5′-ACAAAAAATCAAAAAATCACTTATACAATTCATCCATGCCACC-3′ from the plasmid pCFJ90. The primers 5′-TGATTTTTTGATTTTTTGTTGATTT-3′ and 5′-TTCAAAGAAATCGCCGACTTCAATCGCTCTCAACGTTTCTG-3′ were used to generate the 3′ UTR region of *susi-2*. The vector fragment was PCR amplified with the primers 5′-AGAAACGTTGAGAGCGATTGGTGAGTTCCAATTGATAATTGTGAT-3′ and 5′-GTATTTTGGGAATTGACAGGG-3′ from plasmid pSG274. These five fragments were joined together by Gibson assembly to form the *susi-2::mCherry* repair plasmid. The transgene was integrated onto the *C. elegans*’ chromosome II via a modified counterselection (cs)-CRISPR method [45].

The sgRNAs used in this study for transgene construction are listed in Supplemental Table S3.

### CRISPR/Cas9-mediated gene deletion

Multiple sgRNAs-guided chromosome deletion was conducted as previously described [46]. To construct sgRNA expression plasmids, the 20 bp *unc-119* sgRNA guide sequence in the pU6::*unc-119* sgRNA(F+E) vector was replaced with different sgRNA guide sequences as described previously. Addgene plasmid #47549 was used to express Cas9 II protein. Plasmid mixtures containing 30 ng/μl of each of the three sgRNA expression vectors, 50 ng/μl Cas9 II expressing plasmid, and 5 ng/μl pCFJ90 were co-injected into YY178: *eri-1(mg366);3xflag::gfp::nrde-3(ggIS1)* animals. The deletion mutants were screened by PCR amplification and confirmed by sequencing. The sgRNAs used in this study are listed in Supplemental Table S3.

### RNAi

RNAi experiments were conducted as previously described [47]. HT115 bacteria expressing the empty vector L4440 (a gift from A. Fire) was used as controls. Bacterial clones expressing dsRNA were obtained from the Ahringer RNAi library and were sequenced to verify their identity. The 18S RNAi #1 clone with dsRNA targeting the 18S rRNA was PCR amplified with the primer pairs 5′-CGCAATTTGCGTCAACTGTGG-3′ and 5′-TCTTCTCGAATCAGTTCAGTCC-3′ from N2 genomic DNA. The L4440 vector fragment was amplified with the primer 5′-ACTGAACTGATTCGAGAAGActtgatatcgaattcctgcagc-3′ and 5′-CACAGTTGACGCAAATTGCGCTTATCGATACCGTCGACCTC-3′. These two fragments were joined together to generate the dsRNA expression plasmid targeting 18S rRNA. The 18S RNAi #2 clone with dsRNA targeting the 18S rRNA was PCR amplified with the primer pairs 5′-TCTATCCGGAAAGGGTGTCTGC-3′ and 5′-CACTCCACCAACTAAGAACGGC-3′ from N2 genomic DNA. The L4440 vector fragment was amplified with the primer 5′-CGTTCTTAGTTGGTGGAGTGcttgatatcgaattcctgcagcc-3′ and 5′-AGACACCCTTTCCGGATAGActtatcgataccgtcgacctcga-3′. These two fragments were joined together to generate the dsRNA expression plasmid targeting 18S rRNA.

### Chromatin immunoprecipitation (ChIP)

ChIP experiments of H3K9me3 or H3K27me3 were performed as previously described with hypochlorite-isolated embryos [20]. Briefly, after crosslinking, samples were sonicated 23 cycles (each cycle: 30 seconds on and 30 seconds off) with a Bioruptor UCD-200 (Diagenode). Lysates were precleared with agarose beads (BBI no. C600957-0005) and then immunoprecipitated with 2 μl anti-trimethylated H3K9 antibody (Millipore no. 07-523) or 2 μl anti-trimethylated H3K27 antibody (Millipore no. 07-449). ChIP signals were normalized to levels of *eft-3* and the data were expressed as ratios of indicated animals exposed to ± dsRNA.

For Pol I transcription, ChIP experiments were performed with hypochlorite-isolated young adults. After cross-linking, samples were resuspended in FA buffer (50 mM Tris/HCl at pH 7.5, 1 mM EDTA, 1% Triton X-100, 0.1% sodium deoxycholate, 150 mM NaCl) containing proteinase inhibitor tablet (Roche, 04693116001) and sonicated for 23 cycles at medium output (each cycle: 30 seconds on and 30 seconds off) with a Bioruptor 200. Lysates were precleared and then immunoprecipitated with 1.5 μL of anti-GFP antibody (Abcam, ab290) for GFP::RPOA-2 overnight at 4°C. Antibody-bound complexes were recovered with Dynabeads Protein A. DNA was treated with RNase (Roche) and Proteinase K (New England Biolabs).

Quantitative real-time PCR (qPCR) was performed using an MyIQ2 machine (Bio-Rad) with SYBR Green Master Mix (Vazyme, Q111-02). The primers used in this work are listed in Table S4.

### Deep sequencing of small RNAs and bioinformatic analysis

Deep sequencing of small RNAs and bioinformatic analysis were conducted as previously described [14]. Briefly, total RNAs were isolated from L3 stage worm using a dounce homogenizer (pestle B) in TRIzol solution (Invitrogen) followed by DNase I digestion (Fermentas, no. 18068015). 3xFLAG::GFP::NRDE-3-associated siRNAs were isolated from L3 stage worm lysates as described previously [5, 21]. The lysate was pre-cleared with protein G-agarose beads (Roche) and incubated with anti-FLAG M2 agarose beads (Sigma #A2220). The beads were washed extensively and were eluted with 100 μg/ml 3xFLAG peptide (Sigma). The eluates were incubated with TRIzol reagent followed by isopropanol precipitation and DNase I digestion (Fermentas). To facilitate 5′-phosphate-independent deep sequencing, the precipitated RNAs were treated with calf intestinal alkaline phosphatase (CIAP, Invitrogen), re-extracted with TRIzol, and treated with T4 polynucleotide kinase (T4 PNK, New England Biolabs) in the presence of 1 mM ATP.

Small RNAs were subjected to deep sequencing using an Illumina platform (Novogene Bioinformatics Technology Co., Ltd). Briefly, small RNAs ranging from 18 to 30 nt were gel-purified and ligated to a 3’ adaptor (5’-pUCGUAUGCCGUCUUCUGCUUGidT-3’; p, phosphate; idT, inverted deoxythymidine) and a 5’ adaptor (5’-GUUCAGAGUUCUACAGUCCGACGAUC-3’). The ligation products were gel-purified, reverse transcribed, and amplified using Illumina’s sRNA primer set (5’-CAAGCAGAAGACGGCATACGA-3’; 5’-AATGATACGGCGACCACCGA-3’). The samples were then sequenced using an Illumina Hiseq platform.

The Illumina-generated raw reads were first filtered to remove adaptors, low quality tags and contaminants to obtain clean reads at Novogene. Clean reads ranging from 18 to 30 nt were mapped to the unmasked *C. elegans* genome and the transcriptome assembly WS243, respectively, using Bowtie2 [48] with default parameters. The number of reads targeting each transcript was counted using custom Perl scripts and displayed by IGV [49]. The number of total reads mapped to the genome minus the number of total reads corresponding to sense rRNA transcripts (5S, 5.8S, 18S, and 26S) and sense protein coding mRNA reads was used as the normalization number to exclude the possible degradation fragments of sense rRNAs and mRNAs.

### Actinomycin D treatment

Actinomycin D (MedChemExpress no. HY-17559, CAS:50-76-0) was prepared to 20 mg/ml in DMSO as stock solution. The actinomycin D stock solution was diluted to 5 μg/ml or 10 μg/ml with concentrated OP50. NGM plates were prepared and placed at room temperature overnight before use. Embryos were placed onto the seeded plates and grown to young adults before collection for ChIP.

### Imaging

Images were collected using Leica DM4 microscopes.

### Statistics

Bar graphs with error bars are presented with mean and standard deviation. All of the experiments were conducted with independent *C. elegans* animals for the indicated N times. Statistical analysis was performed with two-tailed Student’s t-test.

## Data availability

All raw and normalized sequencing data have been deposited to Gene Expression Omnibus under submission number GSE.

## Acknowledgments

We are grateful to the members of the Guang lab for their comments. We are grateful to the International *C. elegans* Gene Knockout Consortium, and the National Bioresource Project for providing the strains. Some strains were provided by the CGC, which is funded by NIH Office of Research Infrastructure Programs (P40 OD010440). This work was supported by grants from the National Key R&D Program of China (2018YFC1004500, 2019YFA0802600, and 2017YFA0102900), the National Natural Science Foundation of China (31671346, 91940303, 31870812, 32070619, 31871300 and 31900434), the China Postdoctoral Science Foundation (2018M632542), Anhui Natural Science Foundation (1808085QC82 and 1908085QC96), the Strategic Priority Research Program of the Chinese Academy of Sciences (XDB39010000), and CAS Interdisciplinary Innovation Team. This study was supported, in part, by the Fundamental Research Funds for the Central Universities.

## Author Contributions

C.Z., S.G. and X.F. designed research; S.L., X.C., T.X., Q.J., Z.X., D.X., and X.Z. performed the research and analyzed data; S.G. and X.F. wrote the paper.

## Supporting online materials

Figs. S1 to S9

Tables S1 to S4

