## Supplemental materials for "Co-surveillance of ribosomal RNA by the exosome complex and nucleolar RNAi in *C. elegans*"

**Supplementary figure legends**

**Fig. S1.** risiRNA accumulates in *dis-3* mutants. (A) Results from the deep sequencing of total small RNAs from indicated animals. Green dashed lines indicate risiRNAs. (B) Schematic of ceDIS-3 (top) and sequence alignments of DIS-3 in different organisms: hs, *homo sapiens*; mm, *Mus musculus*; sp, *Schizosaccharomyces pombe*; dm, *Drosophila melanogaster*; and ce, *C. elegans*. (C) Images of seam cells of *eri-1(mg366);dis-3(ust56);gfp::nrde-3;mCherry::dis-3*. mCherry::DIS3 rescued the cytoplasmic localization of GFP::NRDE-3 in *eri-1* animals. Scale bars, 10 µm. (D) Brood size of *dis-3* mutants grown at 20°C and 25°C, respectively. mean ± s.d.; n > 10 animals.

**Fig. S2.** RRF-1 and RRF-2 are required for risiRNA production. (A) Distribution from the deep sequencing of NRDE-3-associated small RNAs in indicated animals. (B) Results from the deep sequencing of total small RNAs from the indicated animals. Green dashed lines indicate risiRNAs.

**Fig. S3.** Alleles of the exosome subunits used in this study. (A) Brood sizes of indicated animals grown at 20°C. Data are presented as mean ± s.d.; n > 10 animals. (B-J) The *ust* alleles were generated via a dual sgRNA-directed CRISPR/Cas9 gene knockout technology in *C. elegans.* The *tm* alleles and *ok* alleles were acquired from National Bioresource Project and the CGC, respectively.

**Fig. S4.** risiRNA directs NRDE-2 to transcription site of rRNAs。 (A) Brood sizes of wild type animals upon RNAi targeting of *rpoa-2*. Data are presented as mean ± s.d.; n=16 animals. (C) Images of young gravid adult animals expressing GFP::RPOA-2. Scale bars, 25 µm. (D) Images of the embryos expressing GFP::RPOA-2. Scale bars, 25 µm. White arrows indicate nucleoli. Scale bars, 10 µm. (D) Images of *C. elegans* embryos expressing GFP:NRDE-2 (green) and mCherry::RPOA-2 (red) after animals were fed with 18S RNAi #2 clone. Scale bars, 10 µm.

**Fig. S5.** Images of the animals expressing indicated transgenes in the presence of actinomycin D. Shown are embryos (A), soma (B), and germline (C). White arrows indicate nucleoli. Scale bars, 25 μm.

**Fig. S6.** Images of the animals expressing indicated transgenes in the presence of actinomycin D. (A) Images show somatic cells expressing GFP::EXOS-10 in the presence of actinomycin D. The percentage of animals with nucleolar localized GFP::EXOS-10 is indicated (% NCL). The number of scored animals is indicated in parentheses. Scale bars, 10 μm. (B) Images of seam cells expressing GFP::NRDE-3 in the presence of actinomycin D. Scale bars, 5 μm.

**Fig. S7.** Images of the animals expressing the indicated transgenes. (A) L3 animals and (B) germline of young gravid adults. Scale bars, 25 μm.

**Fig. S8.** Subcellular localization of the exosome subunits. (A, B) Images of somatic cells of the animals expressing indicated transgenes. The fluorescence intensity indicated by dashed lines was measured by ImageJ, and the values are shown on the right. Scale bars, 5 μm.

**Fig. S9.** Candidate-based RNAi screening was used to identify genes suppressing risiRNA production. (A) Schematics of the alleles that were generated via dual sgRNA-directed CRISPR/Cas9 gene knockout technology. Each of the alleles contains a frameshift mutation and is likely a null allele.

**Table S1.** Candidate-base RNAi screening for factors affecting the subcellular localization of GFP::EXOS-10. The percentage of animals with nucleolar-localized GFP::EXOS-10 is indicated (% NCL).

**Table S2.** List of strains used in this study.

**Table S3.** Sequences of sgRNAs for CRISPR/Cas9-mediated gene editing.

**Table S4.** Sequences of quantitative real-time PCR primers for ChIP experiments.

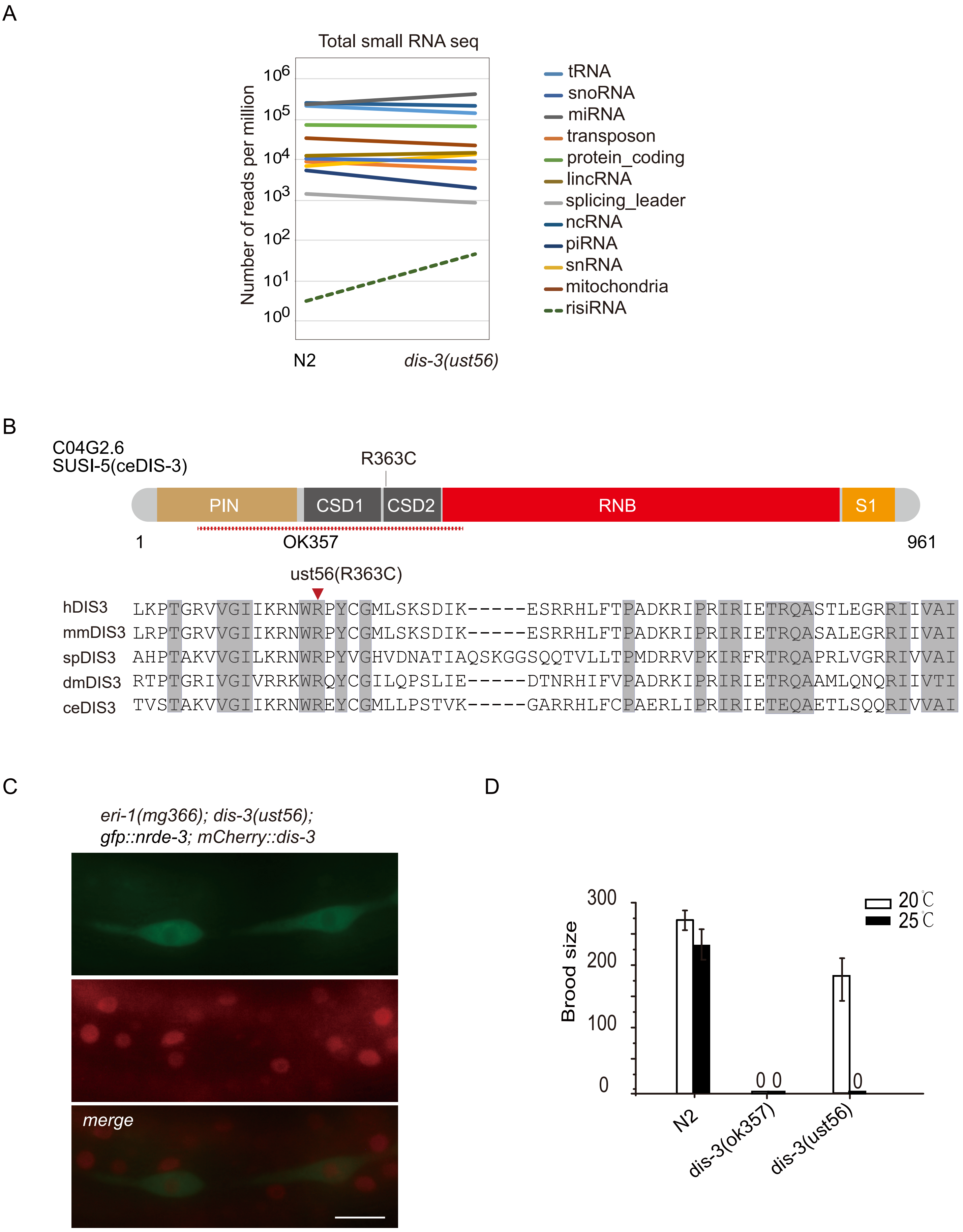
A
Total small RNA seq
Number of reads per million
10^6
10^5
10^4
10^3
10^2
10^1
10^0
N2
dis-3(ust56)
tRNA
snoRNA
miRNA
transposon
protein_coding
lincRNA
splicing_leader
ncRNA
piRNA
snRNA
mitochondria
risiRNA
B
C04G2.6
SUSI-5(ceDIS-3)
R363C
PIN
CSD1
CSD2
RNB
S1
1
OK357
961
ust56(R363C)
hDIS3 LKPTGRVVGIIKRNWRPYCGMLSKSDIK-----ESRRHLFTPADKRIPRIRIETRQASTLEGRRIIVAI
mmDIS3 LRPTGRVVGIIKRNWRPYCGMLSKSDIK-----ESRRHLFTPADKRIPRIRIETRQASALEGRRIIVAI
spDIS3 AHPTAKVVGILKRNWRPYVGHVDNATIAQSKGGSQQTVLLTPMDRRVPKIRFRTRQAPRLVGRRIVVAI
dmDIS3 RTPTGRIVGIVRRKWRQYCGILQPSLIE-----DTNRHIFVPADRKIPRIRIETRQAAMLQNQRIIVTI
ceDIS3 TVSTAKVVGIIKRNWREYCGMLLPSTVK-----GARRHLFCPAERLIPRIRIETEQAETLSQQRIVVAI
C
eri-1(mg366); dis-3(ust56);
gfp::nrde-3; mCherry::dis-3
merge
D
Brood size
300
200
100
0
20℃
25℃
0 0
0
N2
dis-3(ok357)
dis-3(ust56)

Figure S1

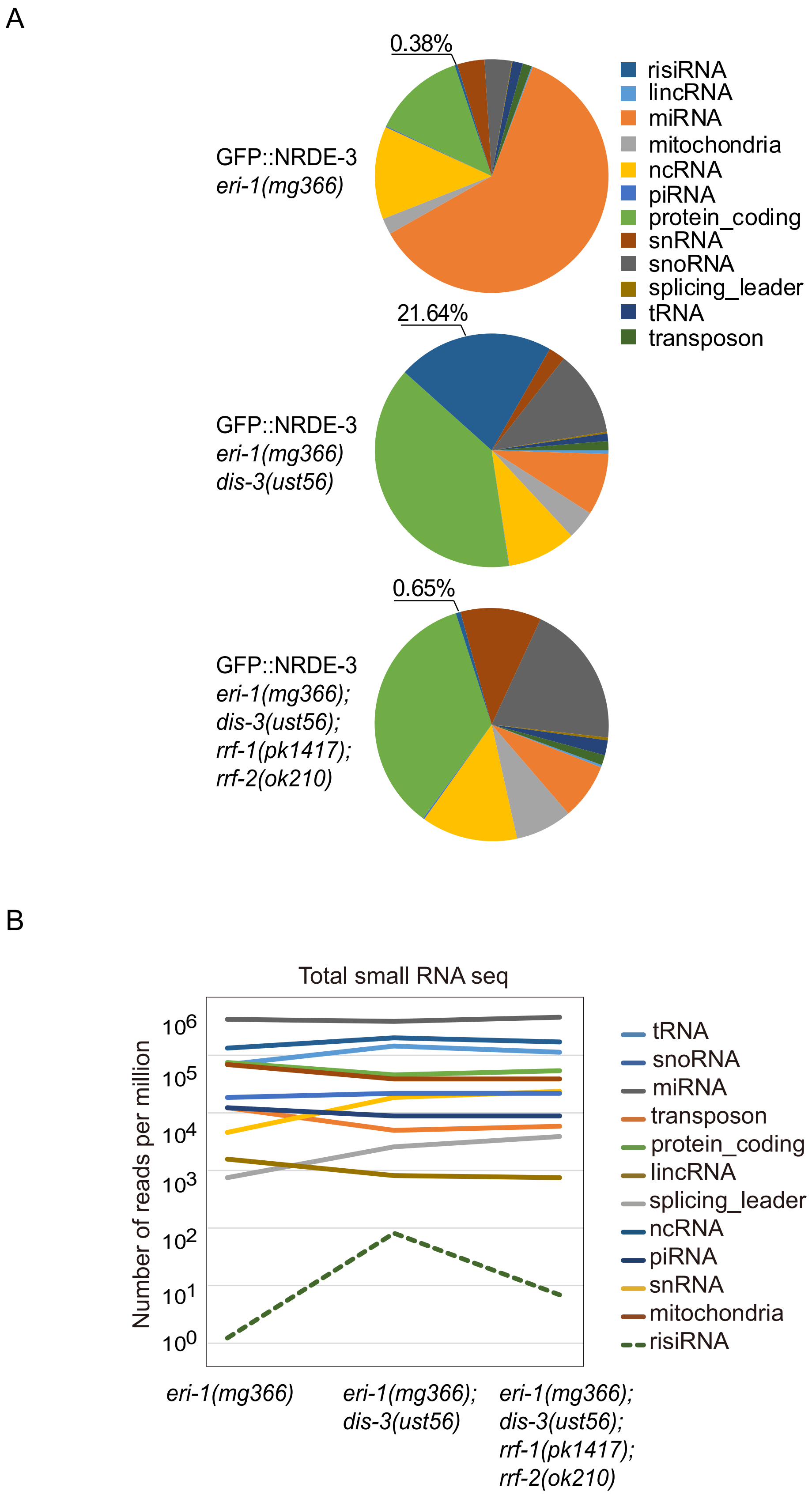
A
0.38%
GFP::NRDE-3
eri-1(mg366)
risiRNA
lincRNA
miRNA
mitochondria
ncRNA
piRNA
protein_coding
snRNA
snoRNA
splicing_leader
tRNA
transposon
21.64%
GFP::NRDE-3
eri-1(mg366)
dis-3(ust56)
0.65%
GFP::NRDE-3
eri-1(mg366);
dis-3(ust56);
rrf-1(pk1417);
rrf-2(ok210)
B
Total small RNA seq
Number of reads per million
10^6
10^5
10^4
10^3
10^2
10^1
10^0
tRNA
snoRNA
miRNA
transposon
protein_coding
lincRNA
splicing_leader
ncRNA
piRNA
snRNA
mitochondria
risiRNA
eri-1(mg366)
eri-1(mg366); dis-3(ust56)
eri-1(mg366); dis-3(ust56); rrf-1(pk1417); rrf-2(ok210)

Figure S2

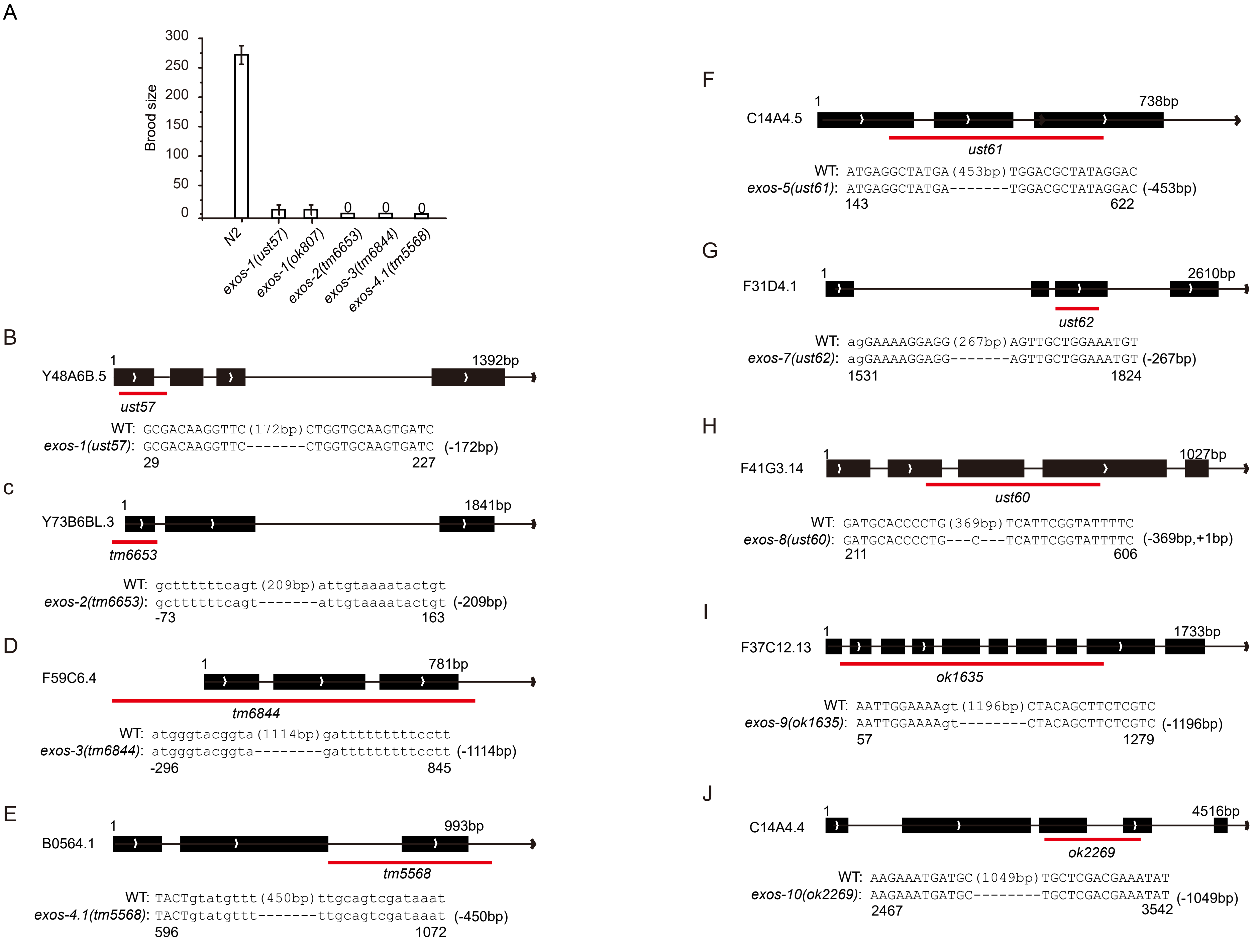
A
300
250
200
150
100
50
0
Brood size
0
0
0
N2
exos-1(ust57)
exos-1(ok807)
exos-2(tm6653)
exos-3(tm6844)
exos-4.1(tm5568)
B
Y48A6B.5
1
1392bp
ust57
WT: GCGACAAGGTTC(172bp)CTGGTGCAAGTGATC
exos-1(ust57): GCGACAAGGTTC-------CTGGTGCAAGTGATC (-172bp)
29
227
C
Y73B6BL.3
1
1841bp
tm6653
WT: gcttttttcagt(209bp)attgtaaaatactgt
exos-2(tm6653): gcttttttcagt-------attgtaaaatactgt (-209bp)
-73
163
D
F59C6.4
1
781bp
tm6844
WT: atgggtacggta(1114bp)gatttttttttcctt
exos-3(tm6844): atgggtacggta--------gatttttttttcctt (-1114bp)
-296
845
E
B0564.1
1
993bp
tm5568
WT: TACTgtatgttt(450bp)ttgcagtcgataaat
exos-4.1(tm5568): TACTgtatgttt-------ttgcagtcgataaat (-450bp)
596
1072
F
C14A4.5
1
738bp
ust61
WT: ATGAGGCTATGA(453bp)TGGACGCTATAGGAC
exos-5(ust61): ATGAGGCTATGA-------TGGACGCTATAGGAC (-453bp)
143
622
G
F31D4.1
1
2610bp
ust62
WT: agGAAAAGGAGG(267bp)AGTTGCTGGAAATGT
exos-7(ust62): agGAAAAGGAGG-------AGTTGCTGGAAATGT (-267bp)
1531
1824
H
F41G3.14
1
1027bp
ust60
WT: GATGCACCCCTG(369bp)TCATTCGGTATTTTC
exos-8(ust60): GATGCACCCCTG---C---TCATTCGGTATTTTC (-369bp,+1bp)
211
606
I
F37C12.13
1
1733bp
ok1635
WT: AATTGGAAAAgt(1196bp)CTACAGCTTCTCGTC
exos-9(ok1635): AATTGGAAAAgt--------CTACAGCTTCTCGTC (-1196bp)
57
1279
J
C14A4.4
1
4516bp
ok2269
WT: AAGAAATGATGC(1049bp)TGCTCGACGAAATAT
exos-10(ok2269): AAGAAATGATGC--------TGCTCGACGAAATAT (-1049bp)
2467
3542

Figure S3

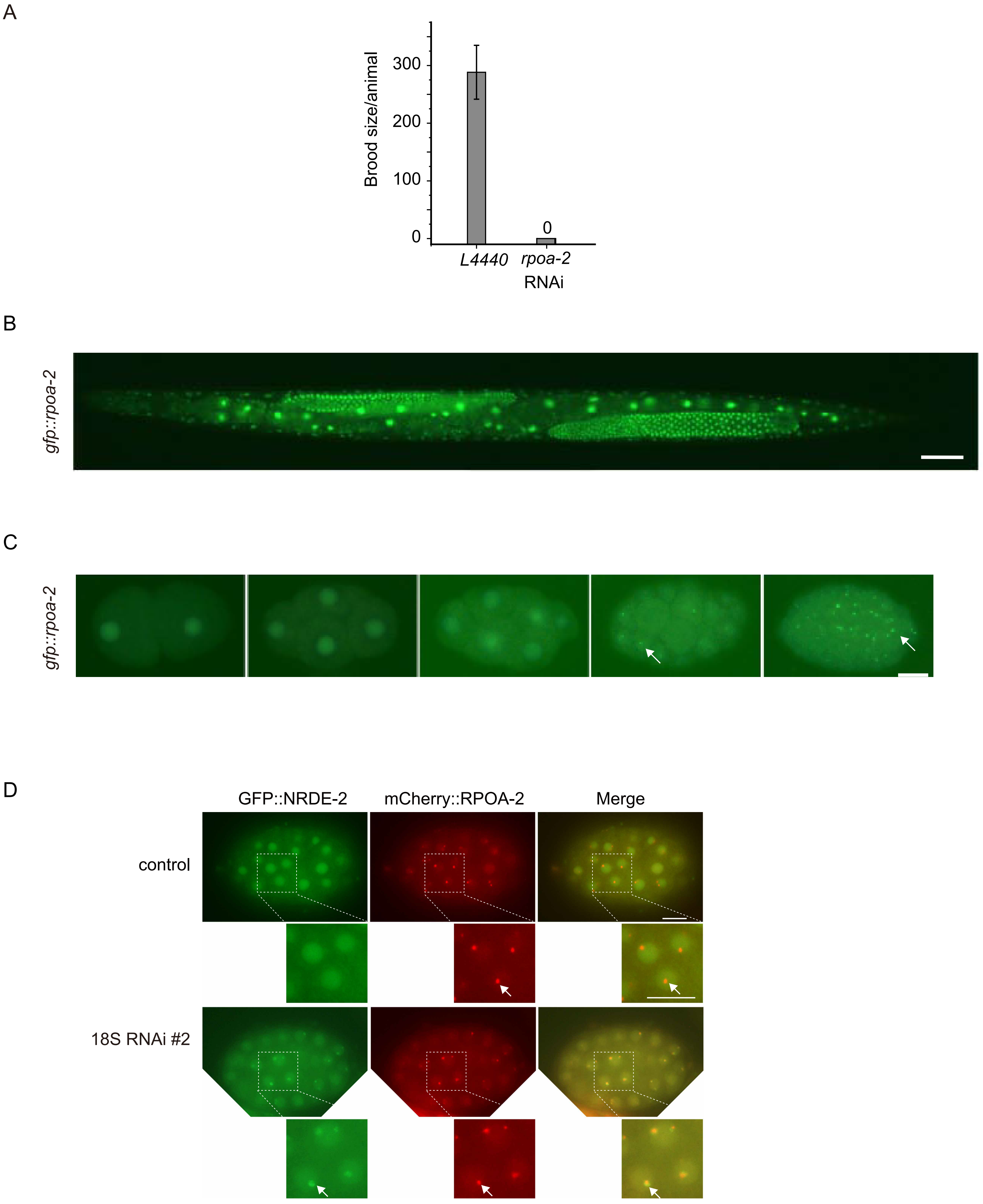
A
Brood size/animal
300
200
100
0
0
L4440
rpoa-2
RNAi
B
gfp::rpoa-2
C
gfp::rpoa-2
D
GFP::NRDE-2
mCherry::RPOA-2
Merge
control
18S RNAi #2

Figure S4

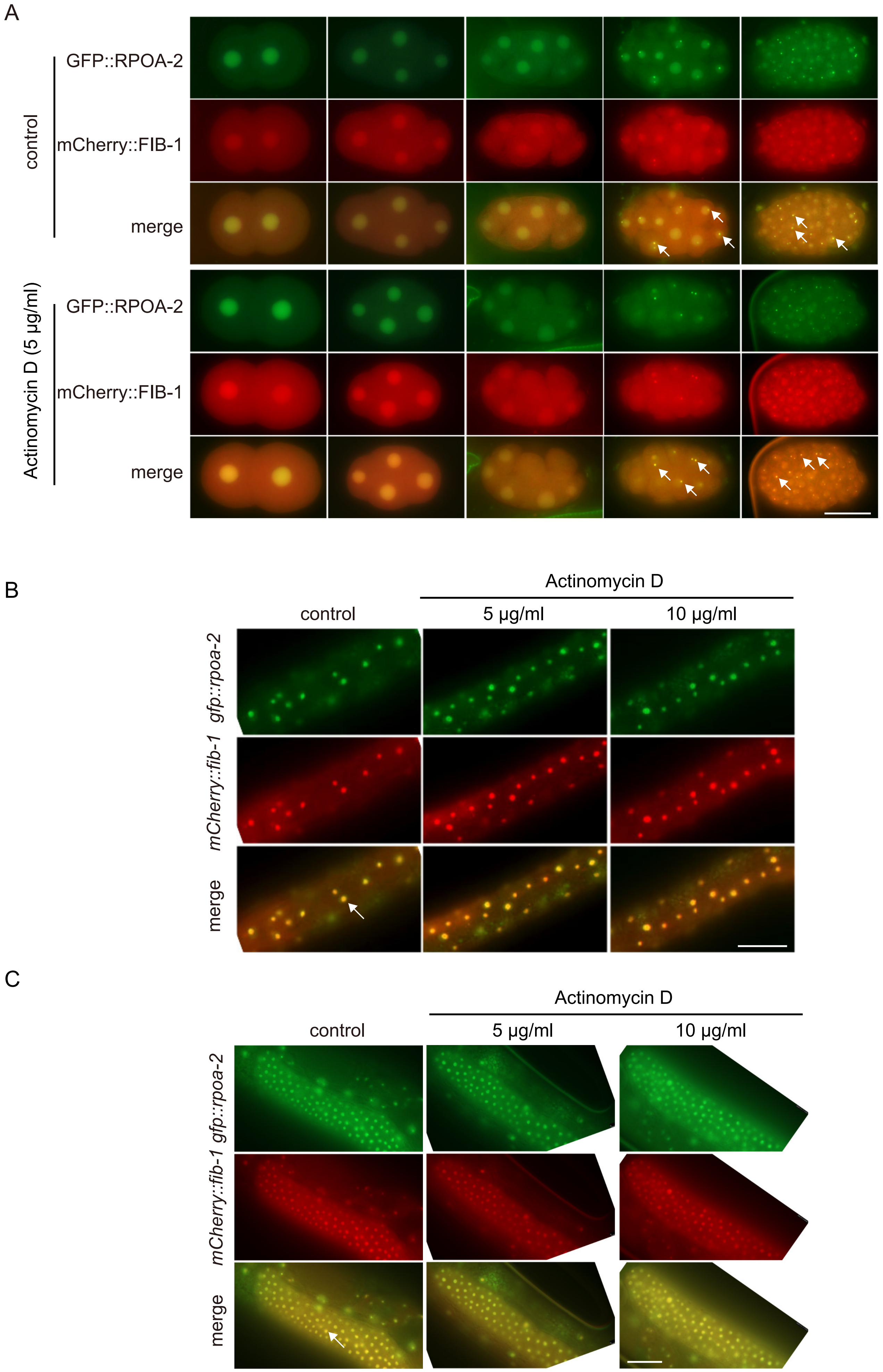
A
control
GFP::RPOA-2
mCherry::FIB-1
merge
Actinomycin D (5 µg/ml)
GFP::RPOA-2
mCherry::FIB-1
merge
B
Actinomycin D
control
5 µg/ml
10 µg/ml
gfp::rpoa-2
mCherry::fib-1
merge
C
Actinomycin D
control
5 µg/ml
10 µg/ml
gfp::rpoa-2
mCherry::fib-1
merge

Figure S5

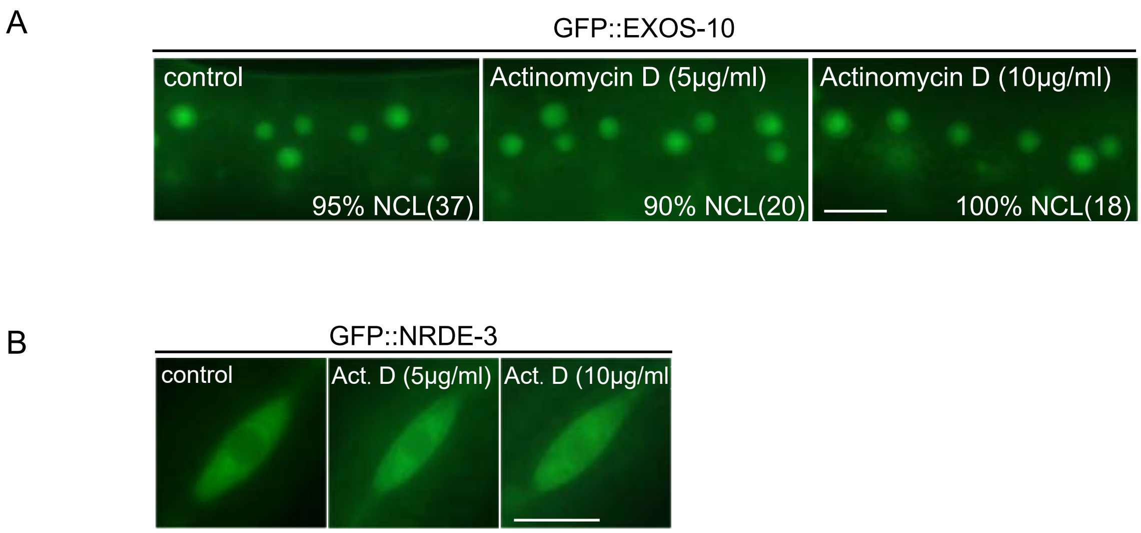
A
GFP::EXOS-10
control
Actinomycin D (5μg/ml)
Actinomycin D (10μg/ml)
95% NCL(37)
90% NCL(20)
100% NCL(18)
B
GFP::NRDE-3
control
Act. D (5μg/ml)
Act. D (10μg/ml

Figure S6

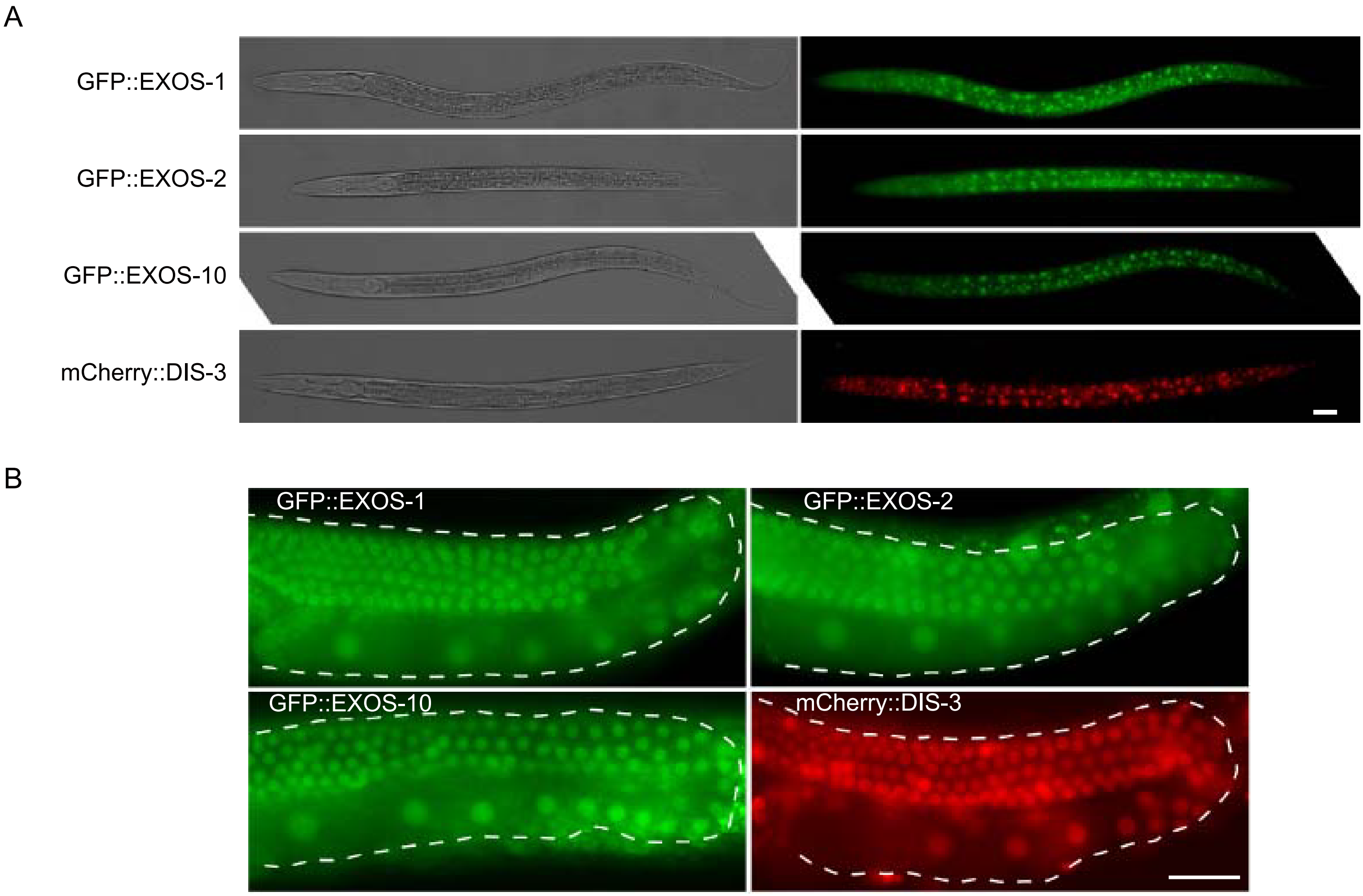
A
GFP::EXOS-1
GFP::EXOS-2
GFP::EXOS-10
mCherry::DIS-3
B
GFP::EXOS-1
GFP::EXOS-2
GFP::EXOS-10
mCherry::DIS-3

Figure S7

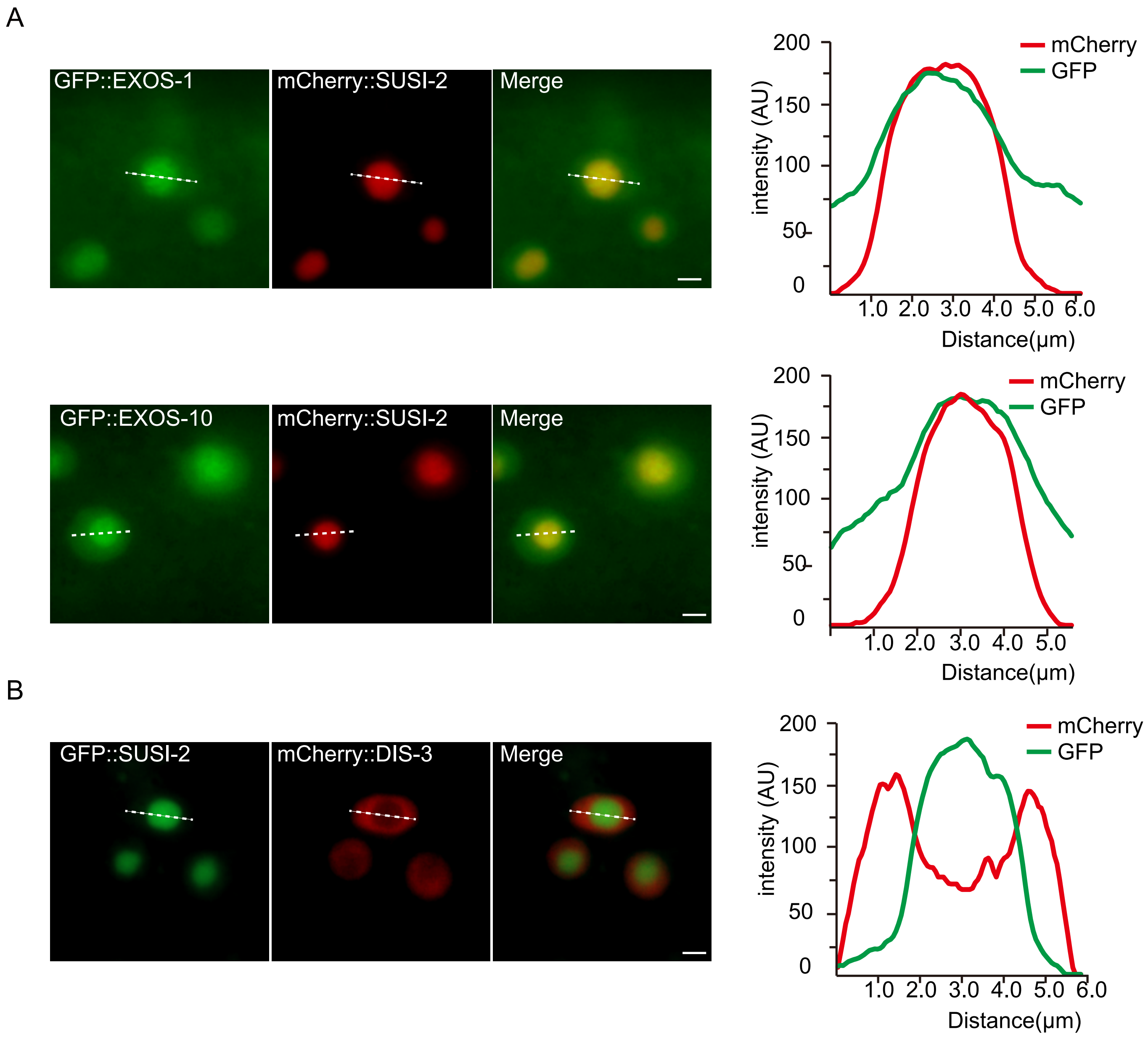
A
GFP::EXOS-1
mCherry::SUSI-2
Merge
mCherry
GFP
intensity (AU)
Distance(μm)
GFP::EXOS-10
mCherry::SUSI-2
Merge
mCherry
GFP
intensity (AU)
Distance(μm)
B
GFP::SUSI-2
mCherry::DIS-3
Merge
mCherry
GFP
intensity (AU)
Distance(μm)

Figure S8

A

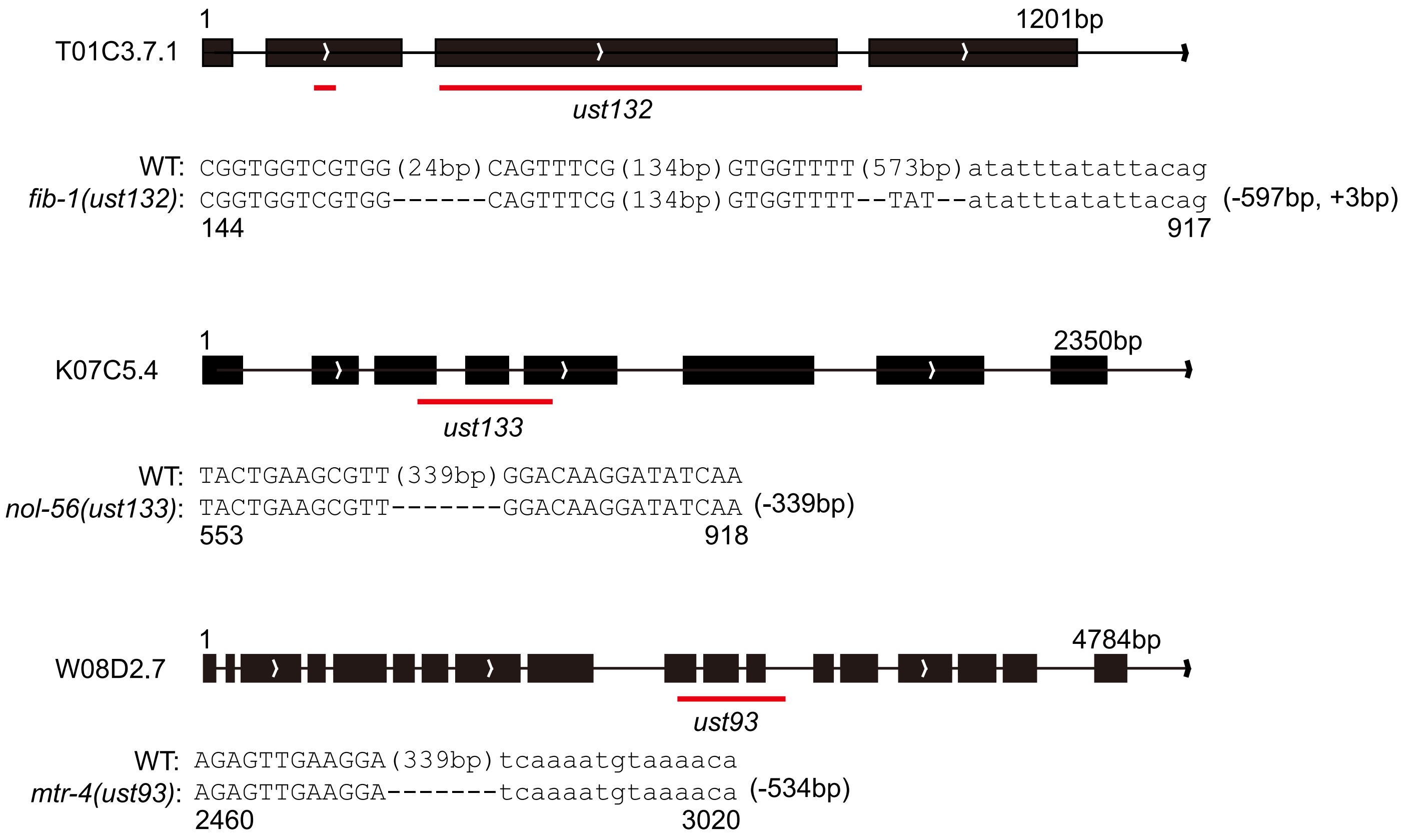

1
1201bp
T01C3.7.1
ust132
WT: CGGTGGTCGTGG(24bp)CAGTTTCG(134bp)GTGGTTTT(573bp)atatttatattacag
fib-1(ust132): CGGTGGTCGTGG------CAGTTTCG(134bp)GTGGTTTT--TAT--atatttatattacag (-597bp, +3bp)
144
917
1
2350bp
K07C5.4
ust133
WT: TACTGAAGCGTT(339bp)GGACAAGGATATCAA
nol-56(ust133): TACTGAAGCGTT-------GGACAAGGATATCAA (-339bp)
553
918
1
4784bp
W08D2.7
ust93
WT: AGAGTTGAAGGA(339bp)tcaaaatgtaaaaca
mtr-4(ust93): AGAGTTGAAGGA-------tcaaaatgtaaaaca (-534bp)
2460
3020


Figure S9

Table S1: Candidate-base RNAi screening for factors affecting the subcellular localization of GFP::EXOS-10.

| gene ID | gene name | NCL% | yeast | human | predicted functions |
|---|---|---|---|---|---|
| L4440(control) | / | 95.5% | | | |
| K07C5.4 | *nol-56* | 0% | NOP56 | NOP56 | histone methyltransferase binding activity and snoRNA binding activity |
| T01C3.7 | *fib-1* | 4.6% | FIB1 | FBL | have RNA binding activity and methyltransferase activity |
| M28.5 | *phi-9* | 11.9% | SNU13 | SNU13 | box C/D snoRNP complex |
| W08D2.7 | *mtr-4* | 22.7% | MTR4 | MTR4 | have ATP binding activity, RNA binding activity, and RNA helicase activity |
| F10B5.1 | *rpl-10* | 31.8% | RPL10 | RPL10 | ribosomal large subunit assembly |
| F37C12.13 | *exos-9* | 60.0% | RRP45 | EXOSC9 | exonucleolytic trimming to generate mature 3'-end of 5.8S rRNA from tricistronic rRNA transcript |
| C04G2.6 | *dis-3* | 68.2% | RRP44 | DIS3 | exosome endoribonuclease and 3'-5' exoribonuclease |
| ZK265.6 | *nol-16* | 75.0% | NOP16 | NOP16 | ribosomal large subunit biogenesis |
| K09B11.2 | *nol-9* | 78.8% | GRC3 | NOL9 | Polynucleotide 5'-kinase involved in rRNA processing |
| F25H8.2 | / | 85.0% | NAF1 | NAF1 | box H/ACA snoRNP assembly |
| K01G5.5 | / | 90.9% | CBF5 | DKC1 | box H/ACA snoRNP complex |
| F32E10.1 | *nol-10* | 91.7% | NOL10 | NOL10 | an ortholog of human NOL10 (nucleolar protein 10); exhibits RNA binding activity |
| C27H6.2 | *ruvb-1* | 95.0% | RUVB1 | RUVBL1 | involved in TOR signaling, box C/D snoRNP assembly |
| Y48A6B.3 | / | 95.0% | NHP2 | NHP2 | box H/ACA snoRNP complex |
| Y66H1A.4 | / | 95.5% | GAR1 | GAR1 | box H/ACA snoRNP complex |

Table S2: Strains used in the work.

| Genotype |
| --- |
| *N2* |
| *CB4856* |
| *susi-5(ust56)* |
| *exos-1(ust57)* |
| *eri-1(mg366);FLAG::GFP::NRDE-3(ggIS1)* |
| *eri-1(mg366);susi-5(ust56);FLAG::GFP::NRDE-3(ggIS1)* |
| *eri-1(mg366);dis-3(ok357);FLAG::GFP::NRDE-3(ggIS1)* |
| *eri-1(mg366);susi-5(ust56);mCherry::SUSI-5(ustIS115);FLAG::GFP::NRDE-3(ggIS1)* |
| *eri-1(mg366);susi-5(ust56);rrf-1(pk1417);FLAG::GFP::NRDE-3(ggIS1)* |
| *eri-1(mg366);susi-5(ust56);rrf-2(ok210);FLAG::GFP::NRDE-3(ggIS1)* |
| *eri-1(mg366);susi-5(ust56);rrf-3(pk1426);FLAG::GFP::NRDE-3(ggIS1)* |
| *eri-1(mg366);susi-5(ust56);rrf-1(pk1417);rrf-2(ok210);FLAG::GFP::NRDE-3(ggIS1)* |
| *eri-1(mg366);exos-1(ust57);FLAG::GFP::NRDE-3(ggIS1)* |
| *eri-1(mg366);exos-2(tm6653);FLAG::GFP::NRDE-3(ggIS1)* |
| *eri-1(mg366);exos-3(tm6844);FLAG::GFP::NRDE-3(ggIS1)* |
| *eri-1(mg366);exos-4.1(tm5568);FLAG::GFP::NRDE-3(ggIS1)* |
| *eri-1(mg366);exos-5(ust61);FLAG::GFP::NRDE-3(ggIS1)* |
| *eri-1(mg366);exos-7(ust62);FLAG::GFP::NRDE-3(ggIS1)* |
| *eri-1(mg366);exos-8(ust60);FLAG::GFP::NRDE-3(ggIS1)* |
| *eri-1(mg366);exos-9(ok1635);FLAG::GFP::NRDE-3(ggIS1)* |
| *eri-1(mg366);exos-10(ok2269);FLAG::GFP::NRDE-3(ggIS1)* |
| *FLAG::GFP::EXOS-1(ustIS112)* |
| *FLAG::GFP::EXOS-2(ustIS113)* |
| *FLAG::GFP::EXOS-10(ustIS114)* |
| *mCherry::SUSI-5(ustIS115)* |
| *mCherry::SUSI-2(ustIS75)* |
| *mCherry::SUSI-5(ustIS115);GFP::SUSI-2(ustIS76)* |
| *susi-5(ust56);FLAG::GFP::EXOS-1(ustIS112)* |

*susi-5(ust56);FLAG::GFP::EXOS-10(ustIS114)*

*FLAG::GFP::EXOS-1(ustIS112);mCherry::SUSI-2(ustIS75)*

*FLAG::GFP::EXOS-10(ustIS114);mCherry::SUSI-2(ustIS75)*

*mCherry::SUSI-5(ustIS115);GFP::SUSI-2(ustIS76)*

*eri-1(mg366);fib-1(ust132);FLAG::GFP::NRDE-3(ggIS1)*

*eri-1(mg366);nol-56(ust133);FLAG::GFP::NRDE-3(ggIS1)*

*eri-1(mg366);mtr-4(ust93);FLAG::GFP::NRDE-3(ggIS1)*

*FLAG::GFP::RPOA-2(ustIS116)*

*FLAG::GFP::RPOA-2(ustIS116);mCherry::FIB-1(ustIS36)*

*GFP::NRDE-2(ustIS117);mCherry::RPOA-2(ustIS116)*

*eri-1(mg366);FLAG::GFP::RPOA-2(ustIS116)*

*eri-1(mg366);nrde-2(gg91);FLAG::GFP::RPOA-2(ustIS116)*

Table S3. Sequences of sgRNAs for CRISPR/Cas9-mediated gene editing.

| | |
|---|---|
| rpoa-2_sg#1 | TTCAGTTCGGCCACAATTCG |
| rpoa-2_sg#2 | ATTGTGGCCGAACTGAACAG |
| rpoa-2_sg#3 | GCGACAGCCACTGTTCAGTT |
| exos-1_sg#1 | ACAAGGTTCTCGACGCGAT |
| exos-1_sg#2 | ATCACTTGCACCAGGTTGT |
| exos-1_sg#3 | ATACTGATGATGTCACATT |
| exos-1_sg#4 | CCGACAGCCATCACTTTGG |
| exos-5_sgRNA #1 | GAGTGATGAGGCTATGACTC |
| exos-5_sgRNA #2 | GTACATGGAATTCAGGATGA |
| exos-5_sgRNA #3 | GCATCCAGAAGTGTGTGCGA |
| exos-7_sgRNA #1 | GGCGAATTGACTGCTCGCGT |
| exos-7_sgRNA #2 | GGTGACGTTGCACCAGATGA |
| exos-7_sgRNA #3 | GTTGAGATTGTCAGCAGCAG |
| exos-8_sg#1 | GAGTCTATCCTGACGGACG |
| exos-8_sg#2 | GTTAGTGATGCACCCCTGG |
| exos-8_sg#3 | TGTGGCAAACCTTTGCCTC |
| exos-8_sg#4 | CGTCAATCGATGCATTGTT |
| nrde-2_sgRNA#1 | GGAACAATGTTTCGAGCGTATGG |
| nrde-2_sgRNA#2 | GAAACATTGTTCATTAAGTTTGG |
| fib-1_sgRNA#1 | GCGGTGGTCGTGGAGGATA |
| fib-1_sgRNA#2 | TGTCCGTCGATGACGGAGC |
| fib-1_sgRNA#3 | CCACTTGAGCAGGTAACCC |
| nol-56_sgRNA#1 | TCGATGCCGCTCATGCTGA |
| nol-56_sgRNA#2 | ATCTGAAGCTTGTCTTCGG |
| nol-56_sgRNA#3 | ATTGCCCTTCTCGATCAGT |
| mtr-4_sgRNA#1 | TGAAGGAATGGCTGTTTCA |
| mtr-4_sgRNA#2 | GTACAAAGTACATTGCATT |
| mtr-4_sgRNA#3 | TCGTATATCAATGGGTTAA |
| mtr-4_sgRNA#4 | GCCAAGGCTTTAGCGAATA |

Table S4. Sequences of quantitative real-time PCR primers for ChIP experiments.

| | |
|---|---|
| *eft-3* qRT F | CAAGGATATTCGTCGTGGATCC |
| *eft-3* qRT R | AATCGAGAACTGGAGTGTATCCG |
| *ama-1* qRT F | CGAACCTGCCGATTGATA |
| *ama-1* qRT R | ACCACGATTGACCAACTC |
| *lin-15b* qRT #1F | ACCGAGACCAGCCAATGT |
| *lin-15b* qRT #1R | TCTTCATCCAGTGGTTCATCCT |
| *lin-15b* qRT #2F | CACTGAACTCACAAGACCACAC |
| *lin-15b* qRT #2R | TCCAATTTGAAGTCATCCCTCTG |
| 5ETS-1F | CCTACACTCATGTCTTTGCAGA |
| 5ETS-1R | GCCGTACTATGCAGCAAGG |
| 5ETS-2F | CCACATTCAGAGGCTGGTGA |
| 5ETS-2R | CCTCTCACCAGCCTATCATTCG |
| 5ETS-3F | CGTCTCTCAAATTGCACACTGC |
| 5ETS-3R | CCTGAGACATCACGTCTCAGAC |
| rDNA#1 F | CAAGACCAATACCGCAACATCA |
| rDNA#1 R | TCATTGCGCCGATCCATAGAT |
| rDNA#2 F | GGAGCTAATACATGCAACTATACCC |
| rDNA#2 R | CAGTCGAAACTGACAGTAAACTGC |
| rDNA#3 F | GGATGAGTTATTTCAATGAGTTGAATAC |
| rDNA#3 R | CGCAGCAATAACGAGATACACTAG |
| rDNA#4 F | CTTCGAGTAGCAAGGAGAGG |
| rDNA#4 R | GCACGTAACCTAGATCCAACTAC |
| rDNA#5 F | GGACTGTCGCTTCGAGGTTTAA |
| rDNA#5 R | GCGATGATCCAGCTGCAG |
| rDNA#6 F | GAATCAGTACACTGATTGCCAAAAG |
| rDNA#6 R | GTTCTTCATCGATACTCGATGC |
| rDNA#7 F | CGATAGCGAACAAGTACC |
| rDNA#7 R | TCTCCAAGCAACATCAAC |
| rDNA#8 F | TAATGTCCTCAACCTATTCTCAA |
| rDNA#8 R | GCCAGTTCTGCTTACCAA |
| rDNA#9 F | GGAATCCGACTGTCTAAT |
| rDNA#9 R | CGCTTACTCGAATTACTAC |
| rDNA#10 F | CATACGACTTGGTCTCTTGG |
| rDNA#10 R | CTGCAAAGACATGAGTGTAGG |
| rDNA#11 F | GTCTGGTTTATTCCGATAACGAG |
| rDNA#11 R | GACCTGTTATCGCTCAATCTC |
| rDNA#12 F | CTCGGCTGATCATCAAGACG |
| rDNA#12R | CGATATCAACACACACACGAGAC |
| rDNA#13F | GTGAACTGTCAACGTGAATGC |
| rDNA#13R | GGTCTCATGAGCGAGAAGTTAG |
